# The dynamics and geometry of choice in premotor cortex

**DOI:** 10.1101/2023.07.22.550183

**Authors:** Mikhail Genkin, Krishna V. Shenoy, Chandramouli Chandrasekaran, Tatiana A. Engel

## Abstract

The brain represents sensory variables in the coordinated activity of neural populations, in which tuning curves of single neurons define the geometry of the population code. Whether the same coding principle holds for dynamic cognitive variables remains unknown because internal cognitive processes unfold with a unique time course on single trials observed only in the irregular spiking of heterogeneous neural populations. Here we show the existence of such a population code for the dynamics of choice formation in the primate premotor cortex. We developed an approach to simultaneously infer population dynamics and tuning functions of single neurons to the population state. Applied to spike data recorded during decision-making, our model revealed that populations of neurons encoded the same dynamic variable predicting choices, and heterogeneous firing rates resulted from the diverse tuning of single neurons to this decision variable. The inferred dynamics indicated an attractor mechanism for decision computation. Our results reveal a common geometric principle for neural encoding of sensory and dynamic cognitive variables.

---

Cortical neurons encode external sensory variables and perform internal cognitive computations that transform sensory inputs into motor actions. While these mental processes engage large neural populations, the activity is often coordinated across neurons to form low-dimensional representations on the population level^1^. Such representations were found for sensory and motor variables by mapping out changes in neural activity in response to varying parameters of stimulus or movement^2–5^. For example, the orientation of a visual stimulus is a one-dimensional circular variable encoded in the primary visual cortex, where neural population responses organize on a ring mirroring the topology of the encoded variable^6, 7^ (Fig. 1a). The orientation-tuning curves of single neurons jointly define the embedding shape of this ring in the population state space, that is, the geometry of neural representation^7, 8^ (Fig. 1a).

**Figure 1.**
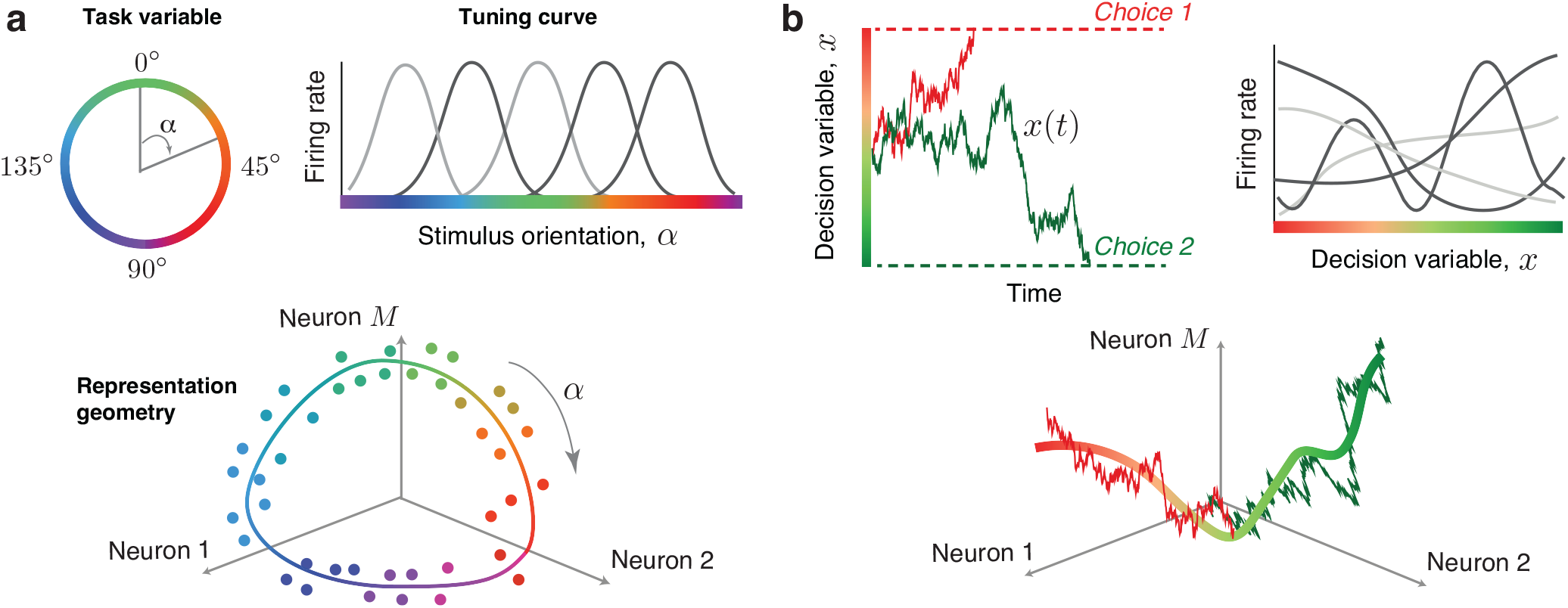
Geometry of neural population codes for sensory and cognitive variables. **a**, Orientation of a visual stimulus is a one-dimensional circular variable *α* (*upper left*). Single neurons in the primary visual cortex encode the orientation of a stimulus with bell-shaped tuning curves, which describe the neuron’s trial-average firing rate as a function of the stimulus orientation (*upper right*). In the population state space, these neural responses form a ring matching the topology of the encoded variable (dots – trial-average population responses to different stimulus orientations indicated by color, the scatter illustrates estimation noise due to a finite number of trials; line – idealized noise-free ring manifold encoding the stimulus orientation). **b**, We hypothesize that the same coding principle holds for dynamic cognitive variables. Specifically, a decision variable *x*(*t*) is a one-dimensional variable representing the dynamics of choice formation on single trials (*upper left*, trajectories colored by the final choice). Single neurons may encode the decision variable with diverse tuning functions, which describe the neuron’s instantaneous firing rate as a function of the decision variable value (*upper right*). During decision formation, neural population responses evolve along a one-dimensional manifold encoding the decision variable in the population state space (*lower panel*, noisy lines illustrate stochastic trajectories of the decision variable on two example trials colored by choice, solid line – idealized noise-free decision manifold). The tuning curves of all neurons jointly define the embedding shape of the decision manifold in the population state space, that is, the geometry of the neural population code for choice.

We hypothesized that neural encoding of dynamic cognitive variables follows the same principle as sensory variables: the encoded variable determines the topology of neural representation, and heterogeneous tuning curves of single neurons define the representation geometry^9^ (Fig. 1b). Testing this hypothesis has been challenging because internal cognitive processes (e.g., decision-making or attention) are not directly observable and unfold with a unique time course on single trials in the sparse and irregular spiking activity of neural populations^10–14^. Thus, these dynamic cognitive representations cannot be revealed by simply averaging neural activity over trials. Moreover, individual neurons show diverse temporal response profiles during cognitive tasks^15–19^, and it is unclear whether single-neuron heterogeneity is compatible with the hypothesis that the neural population represents a low-dimensional cognitive variable^20–22^.

To test our hypothesis, we developed a computational approach for inferring neural population dynamics from spike data that simultaneously learns a model governing the low-dimensional population dynamics on single trials and heterogeneous tuning functions of single neurons to the unobserved population state that define the representation geometry. Two crucial technical advances within this approach make testing our hypothesis possible. First, we perform nonparametric inference over a continuous space of models to discover equations governing population dynamics directly from data^23, 24^, unlike previous methods that tested a small discrete set of models without guarantees that any of these *a priori* chosen models faithfully reflect neural dynamics^11, 21, 25^. Second, our ability to infer heterogeneous tuning functions allows us to reconcile the diversity of single-neuron responses with the population-level encoding of a low-dimensional cognitive variable. In contrast, previous methods assume a rigid monotonic relationship between firing rates of all neurons and latent states and thus capture population dynamics with more latent dimensions, which may not directly correspond to the encoded cognitive variable^11, 21, 25, 26^.

We applied our approach to neural population activity recorded from the primate dorsal premotor cortex (PMd) during decision-making^15^, a cognitive computation described by a decision variable reflecting the dynamics of choice formation on single trials^27, 28^. The neural representation of the decision variable remains unknown since its unique trajectories on single trials are not observable^10, 11^, and decision-related responses of cortical neurons are complex and heterogeneous^15, 21, 22^. Using our computational approach, we provide three lines of evidence for our hypothesis: in dynamics of single neurons, in neural population dynamics, and in their correspondence with animal’s choices.

## Results

### Neural recordings during decision-making

We analyzed spiking activity recorded with linear multielectrode arrays from PMd of two monkeys performing a decision-making task^15^ (Fig. 2a). The monkeys discriminated the dominant color in a static checkerboard stimulus composed of red and green squares and reported their choice by touching the corresponding left or right target when ready (a reaction-time task). We varied the stimulus difficulty across trials by changing the proportion of the same-color squares in the checkerboard and grouped trials into four stimulus conditions according to the response side indicated by the stimulus (left versus right) and stimulus difficulty (easy versus hard, Fig. 2a, Methods).

**Figure 2.**
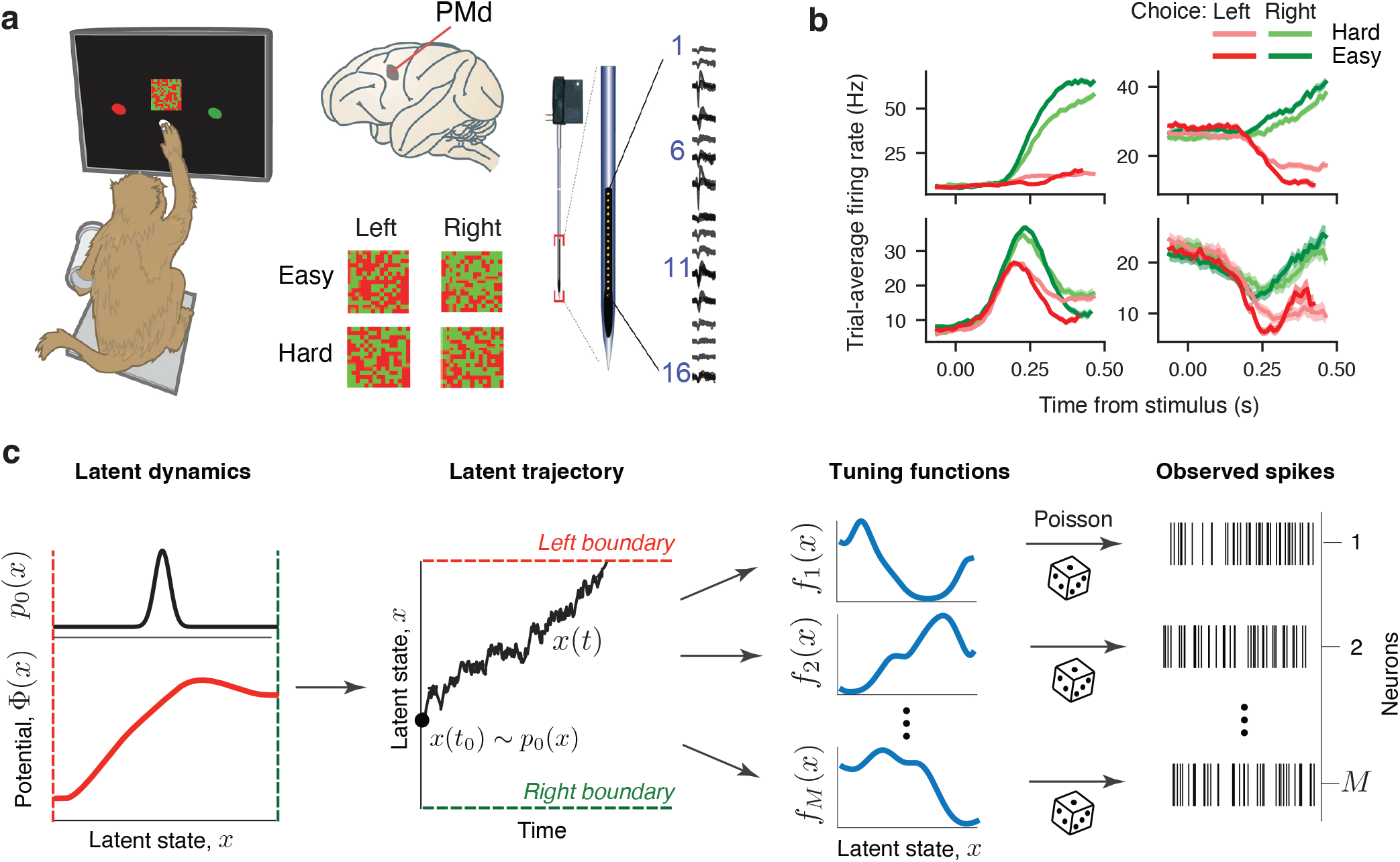
Recording and modeling spiking activity during decision making. **a**, Monkeys discriminated the dominant color in a static checkerboard stimulus composed of red and green squares and reported their choice by touching the corresponding target (*left*). While monkeys performed the task, we recorded spiking activity with 16 channel multi-electrode arrays from PMd (*right*). Trial conditions varied by the response side indicated by the stimulus (left versus right) and stimulus difficulty (easy versus hard, *middle*). **b**, Trial-average firing rates of four example neurons sorted by the chosen side and stimulus difficulty. While some neurons showed canonical ramping responses (*upper panels*), other neurons showed heterogeneous temporal response profiles (*lower panels*). **c**, A framework for simultaneous inference of neural population dynamics and their embedding shape in the population state space. We model neural population dynamics with the latent dynamical system Eq. 1, in which the deterministic flow field arises from a potential Φ(*x*) (*lower left*) and stochasticity is driven by a Gaussian white noise. On each trial, the latent trajectory *x*(*t*) starts at the initial state *x*(*t*_0_) (*middle*, black dot) sampled from the probability density *p*_0_(*x*) (*upper left*). Trial ends when the trajectory reaches one of the decision boundaries corresponding to left and right choice (*middle*, red and green dashed lines). The observed spikes of each neuron follow an inhomogeneous Poisson process with time-varying firing rate that depends on the latent variable *x*(*t*) via neuron-specific tuning functions *f*_*i*_(*x*) (*right*).

Many single neurons in our recordings had decision-related responses with trial-averaged firing rates separating according to the chosen side (Fig. 2b). While some neurons showed canonical firing rates ramping up or down with a slope dependent on the stimulus difficulty, most neurons exhibited heterogeneous temporal response profiles (Fig. 2b), which may seem incompatible with our hypothesis that all these neurons encode the same dynamic decision variable.

### Flexible inference of single-trial neural dynamics

To discover neural representations of the dynamic decision variable, we modeled neural activity as arising from a latent variable *x*(*t*) governed by a general nonlinear dynamical system equation^23, 24^ (Fig. 2c, Methods):

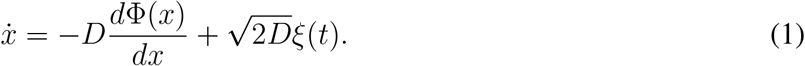

Here Φ(*x*) is a potential function that defines deterministic forces in the latent dynamical system, and *ξ*(*t*) is a Gaussian white noise with magnitude *D* that accounts for stochasticity of latent trajectories. At the beginning of each trial, *x*(*t*) is sampled from the distribution *p*_0_(*x*) of initial states, and the trial terminates when *x*(*t*) reaches one of the decision boundaries in the latent space. We modeled the spikes of each neuron as an inhomogeneous Poisson process with the instantaneous firing rate *λ*(*t*) = *f*_*i*_(*x*(*t*)) changing as a function of the current latent state *x*(*t*). The tuning functions *f*_*i*_(*x*) define the unique dependence of the firing rate on the latent variable *x* for each neuron *i* (Fig. 2c), analogous to tuning curves of single neurons to sensory and motor variables. In our model, Φ(*x*), *p*_0_(*x*), and tuning functions *f*_*i*_(*x*) of all neurons are continuous functions that can take arbitrary nonlinear shapes, which enables us to flexibly discover both the low-dimensional latent dynamics and their nonlinear embedding into the neural population state space.

We simultaneously infer the functions Φ(*x*), *p*_0_(*x*), *f*_*i*_(*x*) and the noise magnitude *D* from spike data by maximizing the model likelihood (Methods, Supplementary Fig. 1, Supplementary Note 2). Thus, our approach discovers equations governing population dynamics directly from data by exploring a continuous space of dynamical system models^23^, each defined by a different potential shape Φ(*x*), of which previously proposed models of decision making^10, 11^ are special cases^24^.

### Decision dynamics in single neurons

First, we examined decision-related dynamics of single neurons by fitting our model to spikes of each neuron separately. We analyzed 128 neurons from monkey T and 88 neurons from monkey O that had sufficiently large choice selectivity and data amount (Methods). Since decision dynamics depend on the stimulus, we first fitted the model for each stimulus condition separately. We noticed that the inferred tuning functions, initial state distribution *p*_0_(*x*) and noise magnitude *D* were similar across four stimulus conditions and only the potential shapes were different (Supplementary Fig. 2). The stability of tuning functions *f*_*i*_(*x*) indicates that stimulus affects only the dynamics of the decision variable but not its encoding in neural activity. In addition, stimulus-independence of *p*_0_(*x*) is expected since stimulus information is not available before stimulus onset. We therefore proceeded with shared optimization in which we fitted the model to all available trials and restricted *f*_*i*_(*x*), *p*_0_(*x*), and *D* to be the same and only allowed the potential Φ(*x*) to differ across stimulus conditions. The shared optimization maximally leverages all available data to produce more reliable and accurate inference (Supplementary Figs. 2-5). The model fit successfully converged for most neurons (monkey T: 117 out of 128 neurons, 91%; monkey O: 67 out of 88 neurons, 76%) which were used in further analyses, showed overfitting for 1 neuron from monkey O (Supplementary Fig. 6, Methods), and showed signs of underfitting for the remaining neurons (monkey T: 11 out of 128 neurons, 9%; monkey O: 20 out of 88 neurons, 23%, Supplementary Fig. 7, a detailed summary of outcomes in Methods).

To examine how well our model accounted for responses of single neurons in our data, we calculated the variance of spike times on single trials explained by the model and compared it to the spike-time variance explained by the baseline prediction based on the trial-average firing rate traces in each condition (Methods). Our model explained significantly more spike-time variance than the baseline (Fig. 3a; monkey T: *p <* 10^−10^, *n* = 111; monkey O: *p <* 10^−10^, *n* = 50, two-sample t-test) and on average accounted for 0.27 ± 0.14 (monkey T) and 0.22 ± 0.13 (monkey O, mean ± std) of the total spike-time variance. Since the total variance of spike times includes an unpredictable point-process variance, we used an independent method to estimate the point-process variance for each neuron^29^ and compared it with the residual variance unexplained by our model (Methods). The point-process variance correlated tightly with the residual variance unexplained by our model (Fig. 3b; monkey T: *r* = 0.80; monkey O: *r* = 0.73, Pearson correlation coefficient), which indicates that our model accounted for nearly all explainable variance in the firing rates of neurons on single trials.

**Figure 3.**
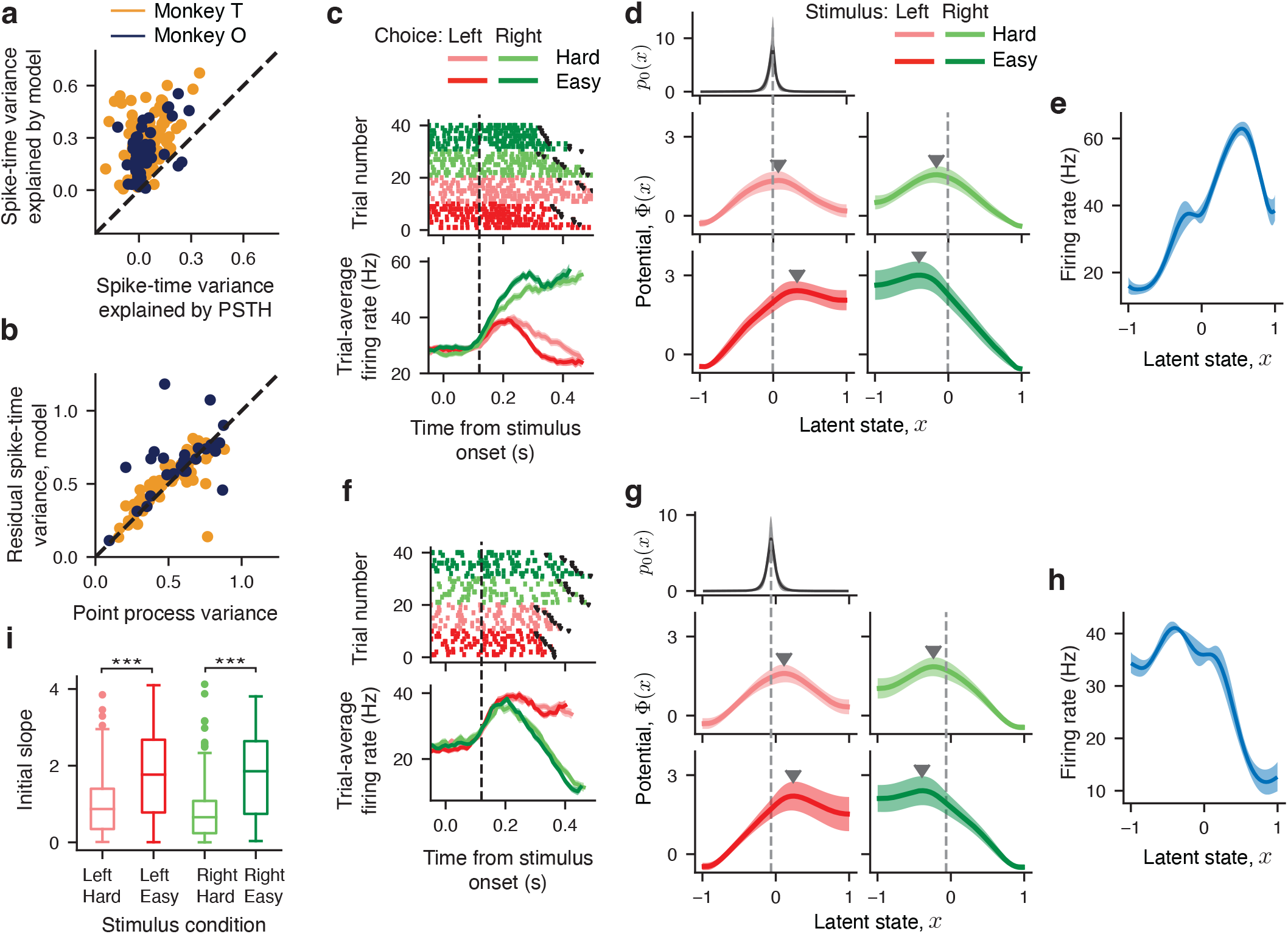
Decision dynamics in single neurons. **a**, Spike-time variance explained by the single-neuron model (y-axis) and by the trial-average firing rate (PSTH, x-axis) for monkey T (orange) and monkey O (purple). Each dot represents data for one neuron averaged across stimulus conditions. **b**, The residual spike-time variance unexplained by the single-neuron model (y-axis) tightly correlates with the point process variance estimated by an independent method (x-axis), which indicates that our model accounts for nearly all explainable firing-rate variance for each neuron. **c**, Spiking activity (*upper panel*) and trial-average firing rates (*lower panel*, PSTH) sorted by the chosen side and stimulus difficulty for an example neuron. Colored dots mark spikes, black dots indicate trial end (reaction time). Error bars are s.e.m. over trials. Time window used for model fitting starts at 120 ms after the stimulus onset (vertical dashed line) and extends until the reaction time on each trial. **d**, Potentials discovered from spikes of the example neuron in c show a single barrier (marked by triangles) in all four stimulus conditions (*middle and lower panels*). The inferred initial state distribution *p*_0_(*x*) shared across conditions (*upper panel*) peaks near the top of the linear slope of the potential on the side corresponding to the correct choice (dashed vertical lines). **e**, The inferred tuning function shared across four stimulus conditions for the example neuron in c. Error bars in d,e are s.t.d. across 10 bootstrap samples. **f**-**h**, Same as c-e for another example neuron. **i**, The distribution of the potential slope at the trial start (at the maximum of *p*_0_(*x*)) for four stimulus conditions. The initial slope is smaller for hard conditions, consistent with longer reaction times than for easy conditions. The box-and-whisker plot shows the initial slope of the potential across all neurons. The center line marks the median, the box extends from the 25th (*Q*_1_) to 75th (*Q*_3_) percentiles, and the whiskers extend from the box to either 1.5× the interquartile range (*Q*_3_ *− Q*_1_) or the most extreme outlier, whichever is closer to the box. The outliers outside of the whiskers are shown with dots.

Our model revealed that despite heterogeneous trial-average responses, single neurons showed remarkably consistent dynamics during choice formation on single trials (Fig. 3c-h), which provides the first line of evidence for our hypothesis. In all stimulus conditions, the inferred potentials displayed the same features: a nearly linear slope towards the decision boundary corresponding to the correct choice and a single potential barrier separating it from the boundary corresponding to the incorrect choice (Fig. 3d,g). The inferred distribution of initial states *p*_0_(*x*) was narrow and centered near the top of the linear slope, indicating that latent trajectories evolve smoothly towards the correct choice but have to overcome the potential barrier towards the incorrect choice (Fig. 3d,g). We observed the same potential shape with a single barrier in all stimulus conditions for the overwhelming majority of single neurons (monkey T: 102 out of 117 neurons, 87%; monkey O: 66 out of 67 single neurons, 98.5%, more examples in Supplementary Fig. 8), and the remaining neurons had a potential with either no barrier or two barriers in at least 1 stimulus condition and a single-barrier potential in the remaining stimulus conditions (a detailed summary of outcomes in Methods). For easy stimulus conditions, the potentials had a higher barrier and steeper slope than for hard conditions (Fig. 3d,g,i; easy vs. hard; left stimulus: *p <* 10^−10^, *n* = 184; right stimulus: *p* =*<* 10^−10^, *n* = 184, two-sample t-test), predicting more latent trajectories reaching the correct choice boundary and faster reaction times which is consistent with the animals’ behavior. The heterogeneity of trial-average responses across neurons (Fig. 3c,f) resulted from the different shapes of the inferred tuning functions (Fig. 3e,h). These results reject the idea that decision-related dynamics are heterogeneous across neurons^21^, which would correspond to diverse shapes of the potential Φ(*x*). Instead, we find that during decision formation, the overwhelming majority of single neurons in PMd follow the same dynamics described by a single-barrier potential and diverse tuning functions account for the heterogeneity of their responses.

### Decision dynamics in neural populations

We found that single neurons consistently showed the same dynamics during decision making, but how are these dynamics organized in the population? One possibility is that the dynamics unfold in unison across all neurons, that is, on each trial, all neurons follow the same latent trajectory *x*(*t*), indicating that the entire population encodes the same latent dynamical variable as we hypothesized (Fig. 1b). Alternatively, individual neurons can follow their unique independent trajectories on single trials even if their dynamics are described by the same potential, in which case different neurons can be at different latent states at the same time, for example, evolving towards opposite choice boundaries on the same trial^11^. To test these possibilities, we examined neural population dynamics by fitting our model to spikes of multiple neurons recorded simultaneously in the same session (Methods). When fitting neural population responses, our model assumes that all neurons share the same latent dynamical variable *x*(*t*) and each neuron has its unique tuning function *f*_*i*_(*x*) to this latent variable (Fig. 2c). Thus, the population model has less freedom to explain neural responses than single-neuron models fitted to spikes of each neuron separately. If single-trial dynamics do not unfold in unison across all neurons, then we expect the population model to fit neural responses worse than single-neuron models.

For each monkey, our data yielded 15 sessions that had at least 3 neurons, with the median number of simultaneously recorded neurons per session 6 for monkey T (range 3 to 19), and 4 for monkey O (range 3 to 7). The model fit successfully converged for most sessions (monkey T: 11 out of 15 sessions, 73%; monkey O: 13 out of 15 sessions, 87%) which were used in further analyses, showed overfitting for 1 session in each monkey (Supplementary Fig. 6, Methods), and showed signs of underfitting for the remaining sessions (monkey T: 3 out of 15 sessions, 20%; monkey O: 1 out of 15 sessions, 7%, Supplementary Fig. 7, a detailed summary of outcomes in Methods).

We compared the performance of the population model and single-neuron model in two ways. First, we compared the variance of spike times on single trials explained by the population model and by the single-neuron models fitted separately to each neuron. The population model explained the same amount of spike-time variance as single-neuron models (Fig. 4a; monkey T: population model 0.27 ± 0.16, single-neuron model 0.29 ± 0.14, mean ± std, *p* = 0.59, two-sample t-test, *n* = 80; monkey O: population model 0.19±0.13, single-neuron model 0.19±0.13, mean ± std, *p* = 0.97, two-sample t-test, *n* = 32), which indicates that single neurons participate in the same shared dynamics unfolding on the population level. Second, we used a more stringent leave-one-neuron-out validation, in which we predict spike times of one neuron from the latent variable *x*(*t*) inferred from spikes of all other neurons in the population on single trials (Methods). We performed this analysis for datasets from monkey T which had sufficiently large number of neurons per session (Methods). The amount of spike-time variance explained by the population model in the leave-one-neuron-out validation was significantly greater than the spike-time variance explained by the baseline prediction based on the neuron’s own trial-average firing rate in each condition (Fig. 4a; monkey T: leave-one-neuron-out 0.15 ± 0.16, baseline 0.04 ± 0.09, mean ± std, *p* = 1.1 · 10^−6^, two-sample t-test, *n* = 80). These results are consistent with our hypothesis that the entire population encodes the same latent dynamical variable on single trials.

**Figure 4.**
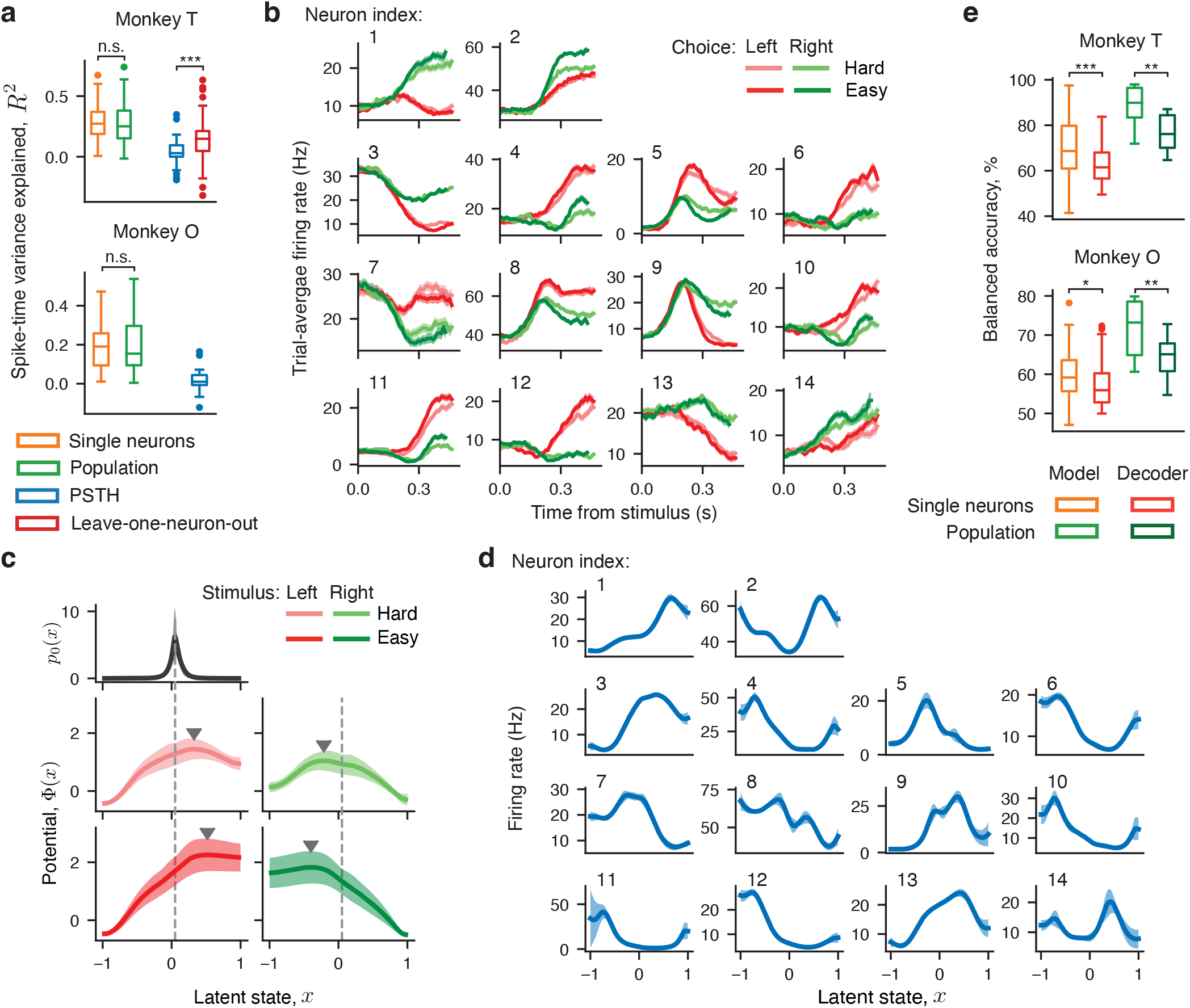
Decision dynamics in neural populations. **a**, The distribution of spike-time variance explained by the population model (green), single-neuron model (orange), trial-average firing rate traces (blue, PSTH), and by the population model in a leave-one-neuron-out validation (red). The box-and-whisker plot shows the explained spike-time variance across all neurons. **b**, Trial-average firing rate traces (PSTHs) sorted by the chosen side and stimulus difficulty for 14 neurons recorded simultaneously on an example session. Error bars are s.e.m. **c**, Potentials governing the population dynamics discovered from spikes of 14 neurons in b show a single barrier (marked by triangles) in all four stimulus conditions (*middle and lower panels*). The inferred initial state distribution *p*_0_(*x*) shared across conditions (*upper panel*) peaks near the top of the linear slope of the potential on the side corresponding to the correct choice (vertical dashed lines). **d**, The inferred tuning functions of 14 neurons in a shared across stimulus conditions. Error bars in c,d are s.t.d. across 10 bootstrap samples. **e**, The distribution of balanced accuracy of predicting monkey’s choice using the single-neuron models (orange), population models (light green), and a logistic regression decoder trained on single-neuron (red) and population activity (dark green). The box-and-whisker plot shows the balanced accuracy of choice prediction across all neurons (for single-neuron models and decoder) and across all sessions (for population models and decoder). The box-and-whisker format is as in Fig. 3i.

Since the population model predicted spike times just as well as single-neuron models, we expect that it recovered the same dynamics as identified in single neurons. Indeed, the dynamics discovered by the population model were consistent with single-neuron results (Fig. 4b-d). For all stimulus conditions, the potential had a linear slope towards the boundary corresponding to the correct choice separated by a potential barrier from the boundary corresponding to the incorrect choice (Fig. 4c). The initial state distribution *p*_0_(*x*) was narrow and centered near the top of the linear slope (Fig. 4c), and the heterogeneity of single-neuron responses (Fig. 4b) was captured in their unique nonlinear tuning functions (Fig. 4d). We observed the same potential shape with a single barrier in all stimulus conditions for the majority of sessions (monkey T: 9 out of 11 sessions, 82%; monkey O: 13 out of 13 sessions, 100%), and the remaining two sessions had a potential with either no barrier or two barriers in at least 1 stimulus condition and a single-barrier potential in the remaining stimulus conditions (detailed summary of outcomes in Methods). The consistency of potential shapes and the high fit quality for the population model provide the second line of evidence for our overall hypothesis.

### Predicting choice from latent dynamics

Our results show that neural populations in PMd encode a one-dimensional dynamic latent variable, and next we tested how this variable was related to the decision-making behavior performed by the animals. We used our models to predict animals’ choices from the neural activity on single trials. On each trial, we decoded the latent trajectory *x*(*t*) from spikes and predicted the choice as the boundary to which this trajectory converged at the reaction time (Methods). Both single-neuron and population models predicted animals’ choices significantly above chance (Fig. 4e; balanced accuracy, mean ± std, one-sample t-test, monkey T: single-neuron model 70 ± 13%, *p <* 10^−10^, *n* = 85, population model 88 ± 7%, *p* = 7 · 10^−8^, *n* = 11; monkey O: single-neuron model 60 ± 7%, *p <* 10^−10^, *n* = 46, population model 72 ± 7%, *p* = 2 · 10^−7^, *n* = 13). This correspondence between the inferred latent variable and animals’ choice is nontrivial, since the information about animals’ choices was not provided to the models during fitting. For a comparison, we trained a logistic regression decoder to predict animals’ choices from spike counts measured in 75 ms sliding window on single trials (Methods). Despite the decoder being directly supervised to predict the choice, our unsupervised models predicted choices with higher accuracy than the decoder (Fig. 4e, monkey T: single-neuron decoder vs. single-neuron models *p* = 2 · 10^−5^, *n* = 85, population decoder vs. population models *p* = 0.006, *n* = 11; monkey O: single-neuron decoder vs. single-neuron models *p* = 0.04, *n* = 46, population decoder vs. population models *p* = 0.006, *n* = 13, two-sample t-test), suggesting that the latent variable inferred by our models is the dynamic decision variable encoded in the spiking activity. Moreover, the population models predicted choices with higher accuracy than single-neuron models (Fig. 4e, monkey T: *p* = 3 · 10^−5^; monkey O: *p* = 7 · 10^−7^, two-sample t-test), reinforcing the conclusion that the decision variable is encoded on the population level. These results provide the third and final line of evidence for our overall hypothesis.

In summary, we find that heterogeneous neural populations in PMd encode the same dynamic decision variable with diverse tuning functions, which define the geometry of the population code for choice formation. This discovery indicates that neural encoding of dynamic cognitive variables and sensory variables follows a common geometric principle.

### Attractor mechanism for decision computation

We discovered that both single-neuron and population activity in PMd are consistently described by the same dynamics—a potential with a single barrier—so what does this result tell us about the mechanism of decision computation in PMd? The dynamics we found in PMd are qualitatively distinct from the stepping and ramping models proposed previously^11, 21, 24, 25^ and, instead, are consistent with the attractor mechanism hypothesized in neural circuit models^30–35^. In the classical discrete attractor spiking network model^30^, two pools of excitatory neurons receive external inputs providing evidence for the left and right choice (Fig. 5a). The excitatory pools compete via a pool of inhibitory neurons that mediates winner-take-all dynamics, such that on each trial, only one excitatory pool elevates the firing rate signaling the network’s choice and the other excitatory pool is suppressed (Fig. 5a).

**Figure 5.**
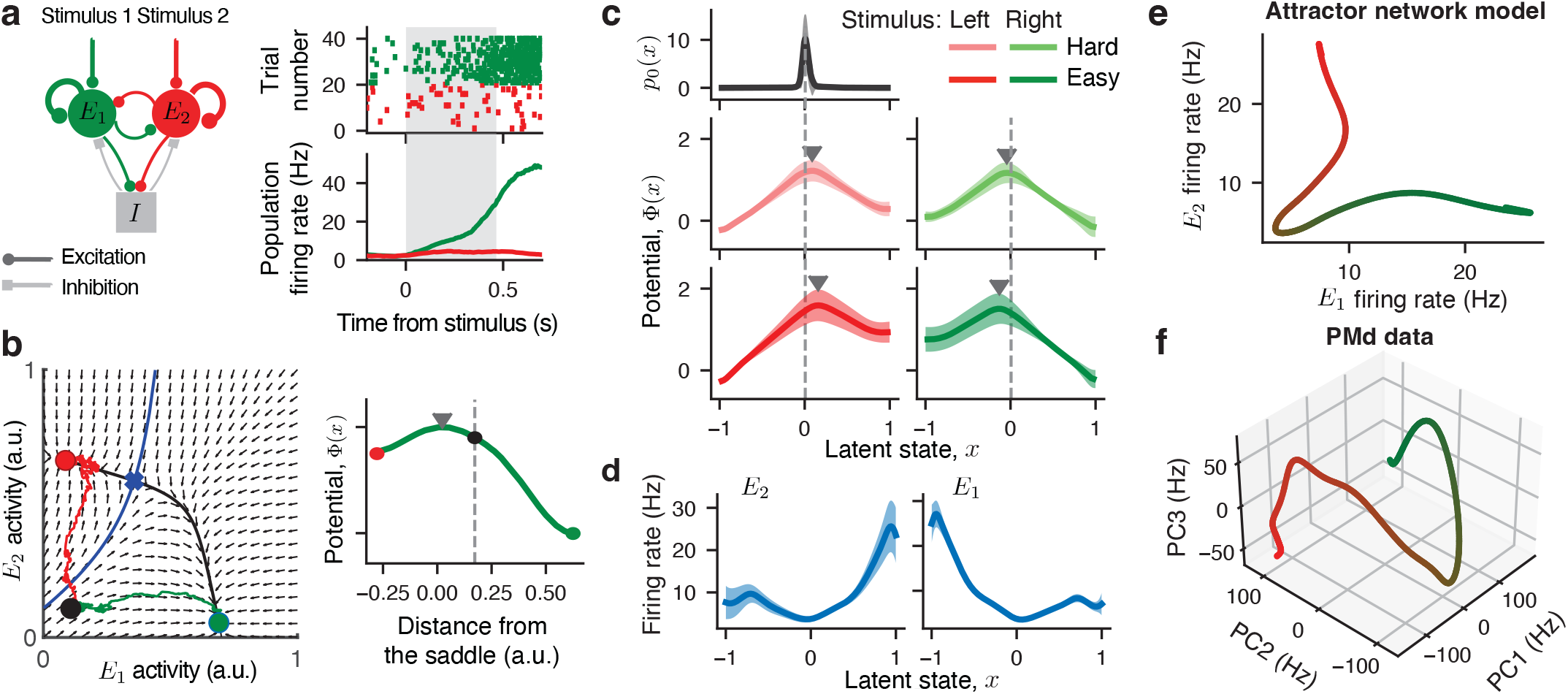
Attractor mechanism for decision computation. **a**, A spiking network model consists of two excitatory neural pools and one inhibitory neural pool (*left*). The recurrent connections are stronger within than across excitatory pools. The inhibitory neurons mediate the winner-take-all competition, in which one excitatory pool elevates the firing rate on each trial signaling the network’s choice and the other excitatory pool is suppressed. The network activity is shown for an example trial: spikes of 40 example neurons (*upper right*) and the population-average firing rates of two excitatory pools (*lower right*) indicated by color. Grey shading indicates time window used for model fitting, starting at the stimulus onset and ending at the reaction time. **b**, Using the mean-field approximation, the network dynamics are reduced to a two-dimensional flow field (arrows) visualized on a phase plane (*left*). On this phase plane, two stable fixed points are attractors corresponding to two choices (red and green circles), which are separated by a saddle point (blue cross). The stable manifold of the saddle point (blue line) is a separatrix dividing the basins of the two choice attractors. At the trial start, the network is initialized in a symmetric low-activity state (black dot). Two example trajectories are shown for a correct (green) and error trial (red). On the error trial, the trajectory has to cross the separatix to reach the incorrect choice attractor. When projected onto a single dimension (the unstable manifold of the saddle point, black line), these dynamics are described by a potential with a single barrier (*right*). **c**, Potentials discovered by fitting spikes generated by the network model show a single barrier (marked by triangles) in all four stimulus conditions, similar to the PMd data. The inferred initial state distribution *p*_0_(*x*) shared across conditions (*upper panel*) peaks near the top of the linear slope on the correct-choice side (vertical dashed lines). **d**, The inferred tuning functions of two example neurons from each excitatory pool. **e**, The decision manifold defined by the tuning functions of neurons in the spiking network model. The manifold is two dimensional because all neurons within each excitatory pool have identical responses and therefore identical tuning functions to the latent population state. **f**, The decision manifold defined by the tuning functions of PMd neurons visualized by projecting on the first three principal components. The decision manifold in PMd has two diverging branches for two choices qualitatively similar to the two-pool network model but with more complex higher-dimensional geometry.

The mechanism of decision computation in this network can be understood using a mean-field approximation that reduces the network to a two-dimensional dynamical system in which the activity of two excitatory pools are the dynamic variables^31, 36^. In the phase plane of this dynamical system, two stable attractors represent two choice alternatives separated by a saddle point (Fig. 5b). The stable manifold of the saddle point is the separatrix, which divides the phase plane into the two attractor basins. When initialized on either side of the separatrix, the network state evolves towards the corresponding attractor following the flow field of the dynamical system. At the trial start, the symmetric low-activity network state falls within the basin of the attractor corresponding to the correct choice. Accordingly, on correct trials, the network’s trajectory smoothly follows the flow field to reach the correct-choice attractor. In contrast, on error trials, the noise drives the trajectory across the separatrix to reach the incorrect-choice attractor, in which case the trajectory moves against the flow field thus overcoming a potential barrier. When projected onto a single dimension (e.g., the unstable manifold of the saddle point), these dynamics are described by a potential that has a linear slope towards the correct choice and a potential barrier separating it from the incorrect choice (Fig. 5b).

To verify this theory, we fitted our population dynamics model to spiking activity generated by simulating the attractor neural network (Methods). From the network activity, the population model inferred potentials with a single barrier in each stimulus condition that were qualitatively similar to the potentials discovered from the PMd data (Fig. 5c, c.f. Fig. 3d,g and Fig. 4c). For all neurons within each excitatory pool, the inferred tuning functions were the same and mirror-symmetric reflections of tuning functions in the opposite pool (Fig. 5d), which is expected in this symmetric network with homogeneous neural pools. Thus, the attractor network and PMd data have similar dynamics but different representation geometry, since PMd neurons had heterogeneous tuning to the dynamic decision variable.

To further compare the representation geometry of the decision variable in the two-pool attractor network and PMd, we visualized the manifold defined by the tuning functions *f*_*i*_(*x*) in the population state space. In the two-pool attractor network, excitatory neurons have only two types of tuning functions and thus naturally form a two-dimensional manifold. We can directly visualize this manifold by plotting the two types of tuning functions against each other (Fig. 5e). The manifold shape closely corresponds to the paths that network trajectories take from the initial state to the choice attractors (Fig. 5e cf. Fig. 5b); this observation reinforces the link between our population dynamics model and the attractor mechanism. Since PMd neurons had heterogeneous tuning functions, the manifold is high-dimensional and cannot be directly visualized. We collected all tuning functions inferred in 24 well-fitted sessions and estimated the manifold dimensionality with principal component analysis (PCA). The first three PCs explained 56.0%, 26.8%, and 5.2% of the total variance, respectively. Projecting the manifold onto the first three PCs revealed a shape with two diverging branches for two choices, similar to the two-pool attractor network but with different higher-dimensional geometry (Fig. 5f). This shape of the decision manifold in PMd could potentially arise from the attractor mechanism distributed across many neurons in a recurrent network with low-dimensional connectivity structure^7, 32, 33, 37^.

## Discussion

We identified the dynamics and geometry of the neural population code for choice formation in the primate premotor cortex. In this code, the instantaneous state of the neural population represents the dynamic decision variable on single trials, and heterogeneous firing rates of single neurons result from their diverse tuning to this decision variable. This discovery was enabled by our nonparametric approach that simultaneously infers neural dynamics and their nonlinear embedding in the neural population state space, and it would not be possible with methods that assume rigid parametric relationships between firing rates and latent state^11, 21, 25, 26^. While previous work measured tuning curves of cortical neurons to the accumulated evidence estimated from behavior^38, 39^, our results suggest that the dynamics of internal cognitive computations may differ across brain areas and thus cannot be described by the same global variable inferred from behavior. Our approach can be applied to discover the dynamics and geometry of other cognitive computations in the brain from neural population activity recordings. The decision dynamics we identified were qualitatively distinct from previously proposed ramping and stepping models^10, 11, 21, 25^ and, instead, suggested an attractor mechanism hypothesized in neural circuit models^30–35^. Thus, our results bridge the gap between the neural manifold and circuit approaches to cognition, motivating future work to understand circuit mechanisms supporting cognitive computations in distributed heterogeneous networks^7^.

## Methods

### Behavioral task and electrophysiological recordings

We analyzed an experimental dataset described previously^15^. Two male monkeys (T and O, *Macaca mulatta*, 6 and 9 years old) were used in the experiments. Experimental procedures were in accordance with NIH Guide for the Care and Use of Laboratory Animals, the Society for Neuroscience Guidelines and Policies, and Stanford University Animal Care and Use Committee (8856).

The monkeys were trained to discriminate the dominant color in a static checkerboard stimulus composed of red and green squares and report their choice by touching the corresponding target. At the start of each trial, a monkey touched a central target and fixated on a cross above the central target. After a short holding period (300 − 485 ms), red and green targets appeared on the left and right sides of the screen. The colors of each side were randomized on each trial. After another short delay (400−1000 ms), the checkerboard stimulus appeared on the screen at the fixation cross and the monkey had to move its hand to the target matching the dominant color in the checkerboard. Monkeys were free to respond when ready. Monkeys were rewarded for the correct choices and received a longer inter trial delays for the incorrect choices. Hand position was monitored by taping an infrared reflective bead to index or middle fingers of each hand and used for measurement of speed and to estimate reaction time.

Difficulty of the task was parameterized by an unsigned stimulus coherence expressed as the absolute difference between the number of red (R) and green (G) squares normalized by the total number of squares |R − G|*/*(R + G). We used a 15 × 15 checkerboard which led to a total of 225 squares. The task was performed with 7 different unsigned coherence levels for monkey T and 8 levels for monkey O. For each stimulus condition, our analysis requires at least a small fraction of incorrect choices so that the neural activity fully explores the decision manifold. Therefore, we only analyzed the 4 most difficult stimulus conditions for each monkey which had sufficient number of error trials. To obtain sufficient data for the model fitting and validation, we merged these four stimulus conditions into two groups combining two easier conditions into one group and two harder conditions into another group. We refer to these two groups as easy and hard stimulus difficulties. Since PMd neurons are selective for the chosen side but not for color^15^, we further divided the trials according to the side indicated by the stimulus (left or right) for each stimulus difficulty (easy or hard), resulting in four analyzed conditions in total.

We recorded neural activity with a linear multi-contact electrode (U-probe) with 16 channels. After online and offline spike sorting (through a combination of MATLAB and plexon offline sorter), the average yield was ∼16 and ∼9 neurons per session for Monkey T and O, respectively, which primarily were well-isolated single units.

### Selection of units for the analyses

After spike sorting and quality control, we had 546 and 450 single neurons and multiunits recorded from monkeys T and O, respectively. From this dataset, we selected units for our analyses based on three criteria: (i) trial-average firing rate traces sorted by the chosen side reach 15 Hz for at least one side at any time between stimulus onset and median reaction time, (ii) the total number of trials across all conditions is at least 560, (iii) selectivity index for the chosen side is greater than 0.6 for monkey T and 0.55 for monkey O. The first two criteria ensure that a unit yields sufficiently large number of spikes for model fitting^23^, and the third criterion selects for units with decision-related activity.

For the first criterion, we used trial-average firing rate traces aligned to stimulus onset (PSTH) sorted by the chosen side, obtained by averaging over trials the spike counts measured in 75 ms bins sliding at 10 ms steps. For the third criterion, we measured the spike count of each neuron on each trial in a [0.2, 0.35] s window aligned to stimulus onset. Selectivity index was defined as the area under the receiver operating characteristic (ROC) curve for discriminating left versus right chosen side based on the spike counts. Selectivity index ranges between 0.5 (no choice selectivity) and 1. For each monkey, we imposed a selectivity index threshold at the median across all neurons (0.6 for monkey T and 0.55 for monkey O), leading to selecting half of all neurons in each monkey. This criterion implies that analyzed neurons had overall lower choice selectivity in monkey O than in monkey T, because choice selectivity was generally lower for neurons from monkey O in our dataset.

128 units for monkey T and 88 units for monkey O passed all three selection criteria and were used in single-neuron analyses. The majority were well-isolated single neurons (monkey T: 127 out of 128 units, 99%, monkey O: 76 out of 88, 86%), and the rest were multiunits. For population analyses, we included sessions that had at least 3 of the selected single units recorded simultaneously, yielding 15 populations for each monkey.

On each trial, we analyzed PMd activity from 120 ms after stimulus onset (the appearance of a checkerboard stimulus on the screen) until the reaction time (the hand leaving the central target), which was estimated at the first time after checkerboard onset when speed of the hand was above 10% of the maximum speed for that trial. The delay of 120 ms was chosen to account for the lag in PMd response to the stimulus. We verified that the model fitting results were the same for a 80 − 120 ms range of delays.

### Inference of latent neural dynamics

We model latent neural dynamics *x*(*t*) as a stochastic nonlinear dynamical system defined by a Langevin equation^24^ (Eq. 1) on the domain *x ∈* [−1; 1]. The deterministic force in this equation arises from the potential Φ(*x*), and *ξ*(*t*) is a white Gaussian noise *(ξ*(*t*)) = 0, *(ξ*(*t*)*ξ*(*t*^*t*^)) = *δ*(*t−t*^*t*^) and *D* is the noise magnitude. At the start of each trial, the initial latent state *x*_0_ is sampled from a distribution with probability density *p*_0_(*x*). We model spikes of each neuron as an inhomogeneous Poisson process with instantaneous firing rate *λ*(*t*) = *f* (*x*(*t*)) that depends on the current latent state via a neuron-specific tuning function *f* (*x*). In population models, all neurons follow the same latent dynamics *x*(*t*) and each neuron *i* has a unique tuning function *f*_*i*_(*x*) (*i* = 1 … *M* where *M* is the number of neurons in the population). In this case, the population dynamics *x*(*t*) are shared by all neurons and the tuning functions *f*_*i*_(*x*) jointly define the nonlinear embedding of these dynamics into the neural population state space. Thus, the model is specified by a set of continuous functions—the potential Φ(*x*), the initial state distribution *p*_0_(*x*), a collection of tuning functions *{f*_*i*_(*x*)*}*—and a scalar noise magnitude *D*. We infer all model components *θ* = *{*Φ(*x*), *p*_0_(*x*), *{f*_*i*_(*x*)*}, D}* from spike data *Y* (*t*).

The spike data consist of multiple trials *Y* (*t*) = *{Y*_*k*_(*t*)*}* (*k* = 1, 2 … *K* where *K* is the number of trials). For independent trials, the total data likelihood is a product of likelihoods of individual trials, we therefore consider here data for a single trial *Y*_*k*_(*t*) and omit the trial index to simplify notation. For each trial, *Y* (*t*) = *{y*_0_, *y*_1_, …, *y*_*N*_, *y*_*E*_*}* is a marked point process, that is, a sequence of discrete observation events. Each observation is a pair *y*_*j*_ = (*t*_*j*_, *i*_*j*_) where *t*_*j*_ is the time of event *j* and *i*_*j*_ is the type of this event. The first and last events mark the trial start time *t*_0_ and trial end time *t*_E_, and the *N* remaining events (*j* = 1, … *N*) are the spike observations where *t*_*j*_ is the time of *j*th spike and *i*_*j*_ is the index of the neuron that emitted this spike. The events are ordered according to their times.

We fit the model by maximizing the data likelihood ℒ [*Y* (*t*)|*θ*] over the space of continuous functions^23, 24^ (Supplementary Note 2.1). The likelihood is a conditional probability density of observing the data *Y* (*t*) given the model *θ* marginalized over all possible latent trajectories:

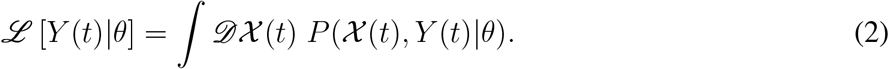

Here *P* (*𝒳* (*t*), *Y* (*t*)|*θ*) is a joint probability density of observing the spike data *Y* (*t*) and a continuous latent trajectory *𝒳* (*t*) given the model *θ*, and the path integral is performed over all possible latent trajectories. To compute the path integral in Eq. 2, we consider a discretized latent trajectory 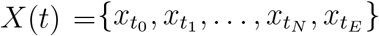, which is a discrete set of points along a continuous path *𝒳* (*t*) at each of the observation times *{t*_0_, *t*_1_, …, *t*_*N*_, *t*_*E*_*}*. Once we calculate the joint probability density *P* (*X*(*t*), *Y* (*t*)) of the discretized trajectory and data, we can obtain the data likelihood by marginalization over all discretized latent trajectories:

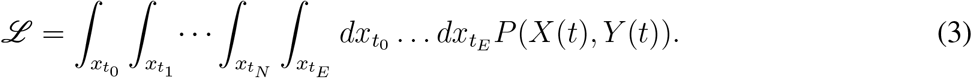

Using the Markov property of the latent Langevin dynamics Eq. 1 and conditional independence of spike observations, the joint probability density *P* (*X*(*t*), *Y* (*t*)) can be factorized^24^:

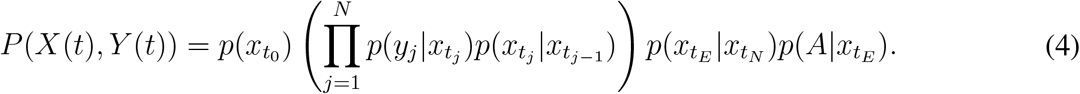

Here 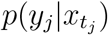 *dt* is the probability of observing a spike from neuron *i*_*j*_ within infinitesimal *dt* of time *t*_*j*_ given the latent state 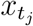 hence 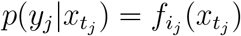 by the definition of the instantaneous Poisson firing rate. 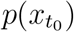 is the probability density of the initial latent state. 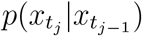 is the transition probability density from 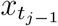 to 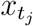 during the time interval between the adjacent spike observations, which accounts for the absence of spikes during this time interval. Finally, the term 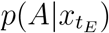 represents the absorption operator, which ensures that only trajectories terminating at one of the domain boundaries at time *t*_*E*_ contribute to the likelihood^24^.

The discretized latent trajectory 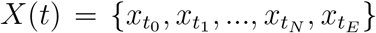 is obtained by marginalizing the continuous trajectory *X* (*t*) over all latent paths connecting 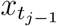 and 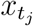 during each interspike interval. These marginalizations are implicit in the transition probability densities 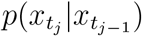 in Eq. 4. The transition probability density 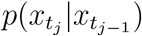 satisfies a modified Fokker-Planck equation which accounts for the drift and diffusion in the latent space and for the absence of spike observations during intervals between adjacent spikes in the data^24^:

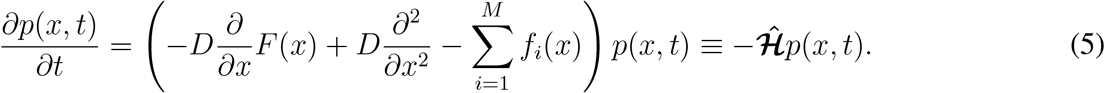

Here *F* (*x*) = −Φ^*t*^(*x*) is the deterministic potential force, and the term 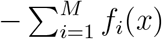 accounts for the probability decay due to spikes emitted by any neuron in the population^24^. The solution of this equation 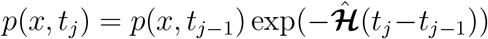 propagates the latent probability density forward in time during each interspike interval. To model the reaction time task, we solve Eq. 5 with absorbing boundary conditions which ensure that trajectories reaching a boundary before the trial end do not contribute to the likelihood^24^. In addition, the absorption operator 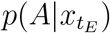 in Eq. 4 enforces that the likelihood includes only trajectories terminating on the boundaries at the trial end time *t*_E_^24^. Together these two conditions ensure that the likelihood includes only trajectories that reach one of the boundaries for the first time at the trial end time.

To fit the model to data, we derived analytical expressions for the gradients of the model likelihood with respect to each of the model components (Supplementary Note 2.2). Instead of directly updating the functions Φ(*x*), *p*_0_(*x*), and *f*_*i*_(*x*) we, respectively, update the force *F* (*x*) = −Φ^*t*^(*x*) and auxiliary functions *F*_0_(*x*) *≡ p*^*t*^_0_(*x*)*/p*_0_(*x*) and 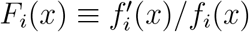. The potential Φ(*x*) is obtained from *F* (*x*) via

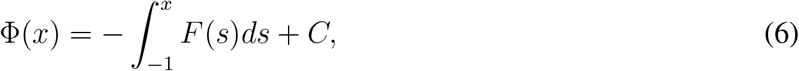

where we fix the integration constant *C* to satisfy ∫_*x*_ exp[−Φ(*x*)]*dx* = 1. The initial state distribution *p*_0_(*x*) is obtained from *F*_0_(*x*) via

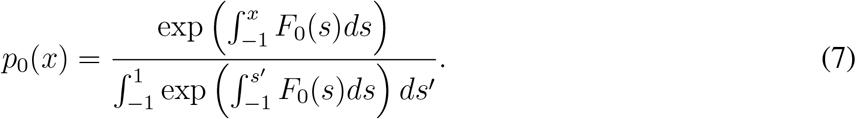

The change of variable from *p*_0_(*x*) to *F*_0_(*x*) allows us to perform an unconstrained optimization of *F*_0_(*x*), while Eq. 7 ensures that *p*_0_(*x*) satisfies the normalization condition for a probability density 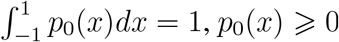. Finally, the tuning function *f*_i_ (*x*) is obtained from *F*_i_ (*x*) via

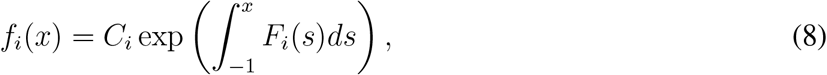

where *C*_*i*_ = *f*_*i*_(−1) is the firing rate at the left domain boundary. This change of variable allows us to perform an unconstrained optimization of *F*_*i*_(*x*), while Eq. 8 ensures the non-negativity of the firing rate *f*_*i*_(*x*) ⩾ 0. We enforce the positiveness of the noise magnitude *D* by rectifying its value after each update *D* = max(*D*, 0), and the same for each constant *C*_*i*_.

We derived analytical expressions for the variational derivatives of the likelihood with respect to each continuous function defining the model *δℒ /δF* (*x*), *δℒ /δF*_0_(*x*), *δℒ /δF*_*i*_(*x*) and the derivatives of the likelihood with respect to scalar parameters *∂ℒ /∂D* and *∂ ℒ /∂C*_*i*_ (Supplementary Note 2.2). We evaluated these analytical expressions numerically for the iterative optimization. To compute the likelihood and its gradients numerically, we use a discrete basis in which all continuous functions, such as *F* (*x*), are represented by vectors, and the transition, emission, and absorption operators are represented by matrices^23, 24^ (Supplementary Note 2.1). Thus, Eq. 4 is evaluated as a chain of matrix-vector multiplications.

### Optimization with ADAM algorithm

We fit the model by minimizing the negative log-likelihood − log *ℒ* [*Y* (*t*)|*θ*] using ADAM algorithm^40^ with custom modifications (Supplementary Note 2.3). The standard ADAM update rule for individual scalar parameters scales their gradients inversely proportional to the average square of their elementwise current and past gradients. Since we optimize over continuous functions *F* (*x*), *F*_0_(*x*), and *{F*_*i*_(*x*)*}*, we scale their gradients by the average function’s *L*^2^-norm defined as ‖*ϕ*(*x*)‖_2_ = ∫_*x*_ |*ϕ*(*x*)|^2^*dx*. We used the following ADAM hyperparameters: *α* = 0.05 for single neurons, *α* = 0.01 − 0.02 for populations, *β*_1_ = 0.9, *β*_2_ = 0.99, = 10^−8^ for both single neurons and population (the definitions of hyperparameters are in Supplementary Note 2.3). We tuned these hyperparameters on synthetic data with known ground-truth. For the scalar parameters *D* and all *{C*_*i*_*}*, we combine ADAM updates with line searches using L-BFGS-B algorithm (L-BFGS-B method from scipy.optimize.minimize toolbox). Since a line search is computationally expensive, we perform only 30 line searches spaced logarithmically over the 5,000 epochs range, such that most line searches are concentrated at early epochs.

We combine ADAM with mini-batch descent randomly splitting the trials from each condition into 20 batches on each epoch. When we perform shared optimization, we fit the model to all available trials restricting *F*_0_(*x*), *{F*_*i*_(*x*)*}*, and *D* to be the same and only allowing the potential force *F* (*x*) to differ across stimulus conditions. In this case, we perform ADAM updates on all batches pooled across four stimulus conditions (80 batches total) in random order on each epoch. We update the force *F* ^*l*^(*x*) that defines the potential in condition *l* only on batches from this condition, and we update all shared components *F*_0_(*x*), *{F*_*i*_(*x*)*}, D, {C*_*i*_*}* on every batch.

We accelerated the optimization algorithm on Graphic Processing Units (GPUs) using cupy library^41^. GPU implementation provides a 5 − 10 fold acceleration over the CPU implementation with the exact factor depending on the spatial resolution of the discrete basis.

### Model selection

ADAM optimization produces a series of models across epochs and we need a model selection procedure for choosing the optimal model. On early epochs, the fitted models miss some true features of the dynamics due to underfitting, whereas on late epochs, the fitted models develop spurious features due to overfitting to noise in the data. The optimal model is discovered on some intermediate epochs. The standard approach for selecting the optimal model is based on optimizing model’s ability to predict new data (i.e. generalization performance), e.g., using likelihood of held-out validation data as a model selection metric^42^. However, optimizing generalization performance cannot reliably identify true features and avoid spurious features when applied to flexible models^23^, which generalize well despite overfitting^43^. We developed an alternative approach for model selection based on directly comparing features of the same complexity discovered from different data samples^23, 24^ (Supplementary Note 2.4). Since true features are the same, whereas noise is different across data samples, the consistency of features inferred from different data samples can separate the true features from noise, and model selection based on feature consistency can reliably identify the correct features^23, 24^.

To compare features discovered from different data samples, we need a metric for feature complexity ℳ. We define feature complexity as the negative entropy of latent trajectories generated by the model^23, 24^ 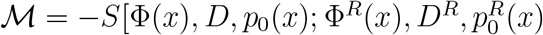. The trajectory entropy^44^ is a functional defined as a negative Kullback-Leibler (KL) divergence between the distributions of trajectories in the model of interest *{*Φ(*x*), *D, p*_0_(*x*)*}* and the distribution of trajectories in the reference model 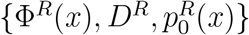. The reference model is a free diffusion in a constant potential (Φ^*R*^(*x*) = const) with the same diffusion coefficient *D* as in the model of interest. We derived the analytical expression for the trajectory entropy for non-stationary Langevin dynamics^24^:

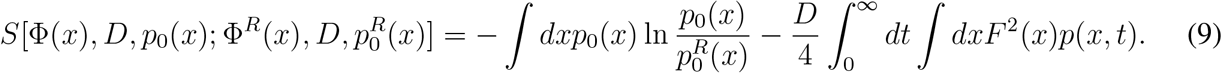

We choose the initial distribution 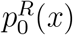 for the reference model to be uniform. We derived an expression for efficient numerical evaluation of Eq. 9 taking the integral over time analytically^24^ (Supplementary Note 2.4). Qualitatively, feature complexity reflects the structure of the potential Φ(*x*): potentials with more structure have higher feature complexity. The reference model with constant potential has zero feature complexity. During model fitting, the feature complexity consistently grows throughout the optimization epochs^24^.

We compare models discovered from two non-intersecting halves of the data *𝒟*_1_ and *𝒟*_2_ to evaluate consistency of their features (Supplementary Fig. 1). We perform the ADAM optimization independently on each data split to obtain two series of models 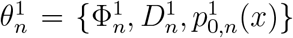 and 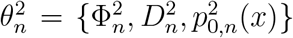 (where *n* = 1, 2 … 5, 000 is the epoch number) fitted on 𝒟_1_ and 𝒟_2_, respectively. We measure feature complexity of these models, 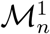 and 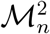, and quantify the consistency of features of the same complexity between models fitted on different data splits. We quantify the consistency of features between two models by evaluating Jensen-Shannon divergence *D*_JS_ between their time-dependent probability densities over the latent space^24^ (Supplementary Note 2.4). At low and moderate feature complexity, the models contain true features of the dynamics in the data and their features agree between data splits reflected in low *D*_JS_ values. At high feature complexity, the models overfit to noise and contain spurious features that do not replicate between data splits, resulting in large *D*_JS_ values. To find the optimal feature complexity, we set the threshold *D*_JS,thres_ = 0.0015 and select ℳ^*∗*^ as the maximum feature complexity for which *D*_JS_ ⩽ (*D*_JS,thres_. This procedure returns two models of roughly the same feature complexity which represent the consistent features of dynamics across data splits. The threshold *D*_JS,thres_ sets the tolerance for mismatch between models and choosing higher *D*_JS,thres_ results in greater discrepancy between models obtained from two data splits. We set *D*_JS,thres_ = 0.0015 based on fitting synthetic data with known ground truth, at this threshold value the selected models reliably matched the ground-truth model.

### Uncertainty quantification

We quantify the estimation uncertainty for fitted models using a bootstrap method^24^. To obtain confidence bounds for the inferred model, we generate ten bootstrap samples by sampling trials randomly with replacement from the set of all trials. To ensure that the two data samples 𝒟_1_ and 𝒟_2_ used for model selection do not overlap, we first randomly split all trials into two equal non-overlapping groups, and then sampled trials randomly with replacement from each group to generate 𝒟_1_ and 𝒟_2_. For shared optimization, we resampled the trials separately for each stimulus condition. For each bootstrap sample, we refit the model and perform model selection using our feature consistency method. We then obtain the confidence bounds for the inferred potential, *p*_0_(*x*) distribution, and tuning functions by computing a pointwise standard deviation across twenty models produced by the model selection on two data splits from each of ten bootstrap samples.

### Outcomes of the model fitting and selection

When fitting our model to spikes of single neurons and populations and performing model selection, we observed three possible outcomes: overfitting, underfitting, and good fit.

In rare cases (monkey T: 0 out of 128 single neurons 0%, 1 out of 15 populations 6.7%; monkey O: 1 out of 88 single neurons 1%; 1 out of 15 populations 6.7%), the model selection produced a model that showed signs of overfitting (Supplementary Fig. 6). We detected overfitting as models with unrealistically high firing rates in the tuning function (hundreds of Hz), disproportionally high noise magnitude (in the range *D ∼* 3 − 5, compared to *D ∼* 0.2 − 0.6 in regular fits) compensated by deep wells in the potential (overall depth of the potential ∼ 20, compared to ∼ 2 in regular fits). These models produced severely underestimated reaction times (reaction time ∼10 ms in the model, compared to ∼500 ms in the data) and did not predict monkey’s choice. This type of overfitting cannot be detected with standard validation approaches^23^, e.g., these models had similar likelihood on training and validation data.

Some selected models showed signs of underfitting in one of two types. In the first type (monkey T: 10 out of 128 single neurons 7.8%, 2 out of 15 populations 13%; monkey O: 11 out of 88 single neurons, 12.5%, 1 out of 15 populations 6.7%), the potentials had the linear slope tilted towards the same boundary in all stimulus conditions, i.e. the model had no decision signal (Supplementary Fig. 7a,b). In the second type (monkey T: 1 out of 128 single neurons 0.8%, 1 out of 15 populations 6.7%; monkey O: 9 out of 88 single neurons, 10%, 0 out of 15 populations 0%), the potentials obtained from two data halves *D*_1_ and *D*_2_ had the linear slope tilted towards the opposite boundaries in at least one stimulus condition (Supplementary Fig. 7c-e). This disagreement about the correct choice side results in *D*_JS_ values rising high early in the optimization, leading to the selection of a model with low feature complexity before all consistent features have been discovered. These both types of underfitting likely arise when a model cannot detect a weak decision signal and mainly fits the condition-independent trend in neural activity.

All remaining models were considered a good fit and were used in further analyses. In these models, we quantified the potential shape by counting the number of barriers in the potential. A barrier is a potential maximum where the force, which is the negative derivative of the potential *F* (*x*) = *−d*Φ(*x*)*/dx*, changes the sign from negative to positive. We also classified a potential minimum next to a boundary as a barrier, because the trajectory must get to the top of the potential to reach the boundary. At a potential minimum, the force changes the sign from positive to negative. We therefore counted the number of sign changes from negative to positive and vice versa in the force *F* (*x*) in each stimulus condition. We used two force functions *F* ^1^(*x*) and *F* ^2^(*x*) produced by the model selection on two data splits *𝒟*_1_ and *𝒟*_2_ (bootstrap samples were not used in this analysis). We counted a sign change to occur within a local region if both *F* ^1^(*x*) and *F* ^2^(*x*) were negative for ten consecutive grid points to the left and positive for ten consecutive grid points to the right of that region, or vice versa. We only counted sign changes that were at least 30 grid points away from the domain boundaries. The overwhelming majority of models had a single-barrier potential in all four stimulus conditions (monkey T: 102 out of 117 single neurons 87%, 9 out of 11 populations 82%; monkey O: 66 out of 67 single neurons, 98.5%, 13 out of 13 populations 100%, Fig. 3e,h, Fig. 4d, Supplementary Figs. 8,9). Some models had a monotonic potential (no barrier) in at least 1 stimulus condition and a single-barrier potential in the remaining conditions (monkey T: 9 out of 117 single neurons 8%, 1 out of 11 populations 9%; monkey O: 0 out of 67 single neurons, 0%, 0 out of 13 populations 0%). The remaining models had a second small barrier in at least 1 stimulus condition and a single-barrier potential in the remaining conditions (monkey T: 6 out of 117 single neurons 5%, 1 out of 11 populations 9%; monkey O: 1 out of 67 single neurons, 1.5%, 0 out of 13 populations 0%). The second barrier was typically shallow and located near the incorrect-choice boundary, where the estimation uncertainty is higher due to lower sampling probability of this region in the data.

We also analyzed the potential shape in models that showed the first type of underfitting with no decision signal. These models had feature complexity similar to good fits, suggesting that the model selection identified similar features in the dynamics. The fit, however, captured only the condition-independent dynamics and missed the weak decision signal. These models can still inform us about the mechanism of decision-making. For example, in the two-pool attractor network model^30^, inhibitory neurons do not have choice selectivity but they still reflect the attractor dynamics with a barrier separating correct and incorrect choices. Many of the models with no decision signal had a single-barrier potential (monkey T: 5 out of 10 single neurons 50%, 0 out of 2 populations 0%; monkey O: 11 out of 11 single neurons, 100%, 1 out of 1 populations 100%), which further supports our finding that the dynamics described by a single-barrier potential were prevalent in our PMd data.

When analyzing spike-time variance explained by our models (Fig. 3a,b, Fig. 4a), for each neuron, we included only stimulus conditions that had at least 600 spikes across all trials. This restriction was necessary for an accurate estimation of the spike-time variance explained, which is computed on raw spike times without binning or smoothing. For single-neuron models, this restriction produced 111 and 50 single neurons for monkey T and O, respectively (Fig. 3a). For population models, this restriction produced 80 and 32 single neurons for monkey T and O, respectively, which were part of the well-fitted populations (Fig. 4a). The comparison between the residual variance unexplained by single-neuron models and the point process variance estimated by the independent method was performed for 64 neurons from monkey T and 27 neurons from monkey O, which had sufficiently high firing rate for the independent method to produce a reliable estimate^29^. For behavior prediction (Fig. 4e), we additionally only included conditions that had at least 5 incorrect choices in both training and validation datasets, which did not change the number of analyzed populations. This condition was necessary for the baseline comparison, which required training a logistic regression decoder for choice prediction. In this analysis, we used all well-fitted population models and the single-neuron models for the exact same set of neurons that were part of the used populations.

### Spike-time variance explained by the models

To quantify how well our models fitted spiking activity on single trials, we computed the amount of variance in raw spike times (without binning or smoothing) explained by a model. We define the spike-time variance using the time rescaling theorem^45^ for doubly stochastic renewal point processes^46, 47^. For a doubly stochastic renewal point process, the total variability in spike times arises from two sources: the variability of the instantaneous firing rate *λ*(*t*) and the variability of the spike-generating point process. The time rescaling theorem states that we can eliminate the firing rate variability by mapping the spike times from the real time *t* to the operational time *t*^*t*^ via squeezing or stretching the time locally in proportion to the cumulative firing rate: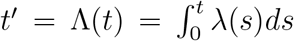. Thus, the variability of rescaled spike times in the operational time reflects only the point process variability. For example, rescaling spike times generated by an inhomogeneous Poisson process yields a homogeneous Poisson process with the firing rate 1 Hz.

We quantify the spike time variability using the squared coefficient of variation of interspike intervals (ISIs) CV^2^, which is the ratio of the ISI variance to the squared mean ISI^48^. The total variance is then 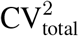 of the raw ISIs calculated in the real time. To compute the variance explained by a model, we use the model to predict the instantaneous firing rate *λ*(*t*) of a neuron on each trial, map the spikes to the operational time using the predicted firing rate *λ*(*t*), compute 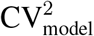 of rescaled ISIs in the operational time, and finally compute the variance explained as:

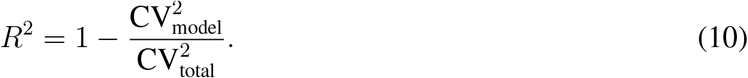

If the predicted instantaneous firing rate *λ*(*t*) faithfully captures the firing-rate dynamics on single trial, then rescaling ISIs eliminates the firing-rate variability leading to 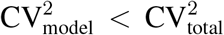. The point process variability still contributes to 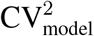, hence *R*^2^ *<* 1 always holds bounded by the unexplainable point process variability. For a Poisson spike-generating process, the rescaled spike times follow a homogeneous Poisson process hence CV^2^ of rescaled ISIs equals one. However, spike statistics of cortical neurons often deviate from the Poisson process^49^, and in our data the residual spike time variance was typically 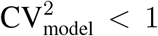 (Fig. 3b) indicating that the spike-generating process is more regular than Poisson^20^. We therefore estimated the point processes variability with an independent method based on more general doubly stochastic renewal point processes^29^ and compared this independently estimated point process variance to the residual variance unexplained by our models. A tight correspondence between the residual spike-time variance unexplained by a model and the independently measured point process variance would indicate that the model accounts for nearly all explainable firing-rate variance in the data.

The independent method for estimating the point process variance is based on doubly stochastic renewal point processes, for which the variance Var(*N*_*T*_) of spike count *N*_*T*_ measured in time bins of size *T* can be partitioned as^29^:

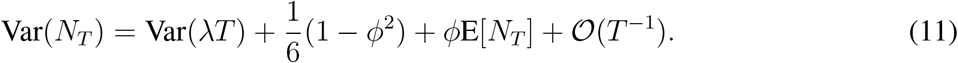

Here *φ* is CV^2^ of ISIs in the operational time, which is a parameter that controls the point process variability, and *λ*(*t*) is the instantaneous firing rate which is assumed to be approximately constant within a single bin. To estimate *φ* from data, we apply Eq. 11 to spike counts measured in two bin sizes *T* and 2*T* to yield two equations, which can be solved to obtain a quadratic equation for *φ*^29^:

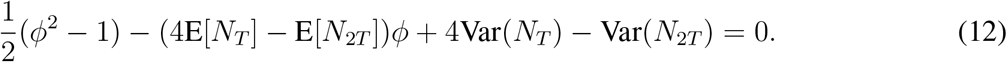

Here, the spike-count mean and variance for each bin size E[*N*_*T*_], E[*N*_2*T*_], Var(*N*_*T*_), Var(*N*_2*T*_) are measured directly from the spike data, and *φ* is the only unknown variable. Thus, we solve Eq. 12 to estimate *φ* from data and compare this *φ* to the residual spike-time variance unexplained by our model.

To predict the instantaneous firing rate *λ*(*t*) with our models, we used the Viterbi algorithm to predict the most probable latent path 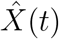 given the observed data *Y* (*t*). We generalized the max-sum Viterbi algorithm with backtracking^50^ to our case of continuous-space continuous-time latent dynamical system (Supplementary Note 2.5). From the most probable latent path 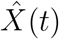, we computed the instantaneous firing rate for a neuron *i* using its tuning function 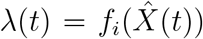. For leave-one-neuron-out validation, we interpolated the instantaneous firing rate with cubic splines to obtain the firing rate at spike times of the left-out neuron. We then rescale spike times via 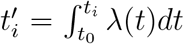 (where *t*_0_ is the trial start time and *t*_1_, …, *t*_*N*_ are the original spike times) using the trapezoidal rule to approximate the time integral and finally compute 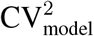 of rescaled ISIs.

We calculated the variance explained by our model using a cross-validation procedure with the same two non-overlapping data splits 𝒟_1_ and 𝒟_2_ as used for the model selection. We used the model fitted on the dataset 𝒟_1_ to predict the instantaneous firing rate and compute *R*^2^ of each neuron on the dataset 𝒟_2_, and vice versa. We analyzed each stimulus condition separately and averaged *R*^2^ across conditions. We report *R*^2^ averaged over the two data splits. For the leave-one-neuron-out validation, we use a population model fitted to spikes of *n* neurons on training trials to predict the instantaneous firing rate of each neuron in turn on validation trials using the latent path predicted with the Viterbi algorithm from the spikes of the remaining *n −* 1 neurons.

As a baseline for the comparison with our models, we also predicted the instantaneous firing rate *λ*(*t*) using the trial-average firing rate traces sorted by the chosen side and stimulus difficulty (i.e. the peristimulus time histogram, PSTH). We computed the trial-average firing rates for the left and right choice trials in a 75 ms window sliding in 10 ms steps on the dataset 𝒟_1_, and used them as a prediction of the instantaneous firing rates for the left and right choice trials on the dataset 𝒟_2_, and vice versa. Thus, this baseline prediction uses the information about the animal’s choice on both the training and validation trials. We analyzed each stimulus condition separately and averaged *R*^2^ across conditions and the two data splits.

### Predicting animal’s choice from neural activity

We used our models to predict animal’s choice from neural activity. We performed a cross-validation procedure with the same two non-overlapping data splits 𝒟_1_ and 𝒟_2_ as used for the model selection. We use the models fitted on the dataset 𝒟_1_ to predict the animal’s choice on the dataset 𝒟_2_, and vice versa, and report the average accuracy over the two data splits. We apply Viterbi algorithm to neural activity on validation trials to predict the most probable latent path 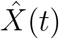. By the design of Viterbi algorithm with absorbing boundary conditions, the trajectory must terminate at one of the domain boundaries, therefore we predict choice as the value of *x*(*t*_*E*_) at the trial end.

As a baseline for the comparison with our models, we also predicted animal’s choice with a logistic regression decoder using the same two data splits for the decoder training and validation. As an input to the decoder, we provide trial-average firing rate traces computed with a 75 ms window sliding in 10 ms steps. We truncated each trial at 0.5 s after stimulus onset resulting in 42-dimensional input vector for single neurons and 42 *× M* -dimensional input vector for populations, where *M* is the number of neurons in the population. We normalize the inputs to have zero mean and unit variance across trials for each condition in each time bin.

Our data is imbalanced as monkeys make more correct choices than errors, especially on easy trials. We therefore report balanced accuracy for both our models and the linear decoder (Fig. 4e). The balanced accuracy is the average between true positive and true negative rates.

### Spiking neural network model

We simulated a spiking recurrent neural network model of decision making with the same parameters as in Ref.^30^ using a python package Brian 2^51^. We only changed the value of the NMDA conductance for inhibitory neurons from *g*_NMDA_ = 0.13 nS to *g*_NMDA_ = 0.128 nS to match the reaction times of the spiking network to the experimental data. We simulated 4 stimulus conditions based on the stimulus difficulty (easy versus hard) and side (left versus right) for comparison with our PMd data. We set the stimulus coherence parameter *c* = 17.5% for easy and *c* = 7.5% for hard stimulus conditions and generated ∼3,200 trials of data per condition. The reaction time was defined on each trial as time when one of the population firing rates (smoothed with a moving average over 200 ms time window) crosses the threshold of 30 Hz. We fitted our population model to responses of two neurons from each of the two selective excitatory pools (i.e. four simultaneous neural responses in total). We performed the same shared optimization across four conditions as for the PMd data using the same hyperparameters for optimization and model selection. To obtain the decision manifold for the network model (Fig. 5e), we plotted the inferred tuning functions of two neurons from different excitatory pools against each other, because tuning functions of all neurons from the same pool are identical.

We used the mean-field approximation to reduce the network’s dynamics to a two-dimensional dynamical system model with the same parameters as in Ref.^31^. To calculate the potential along the unstable manifold of the saddle, we numerically integrated the flux-free flow field along this manifold, 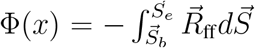. Here 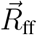 is the flux-free flow field of the reduced two-variable model, and 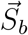 and 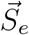 are the start and end points of the manifold, respectively. To find the flux-free flow field, we solved the Poisson equation 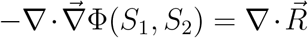, where 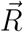 is the full flow field of the reduced two-variable model. The flux-free flow field is then 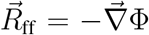. To find the stable and unstable fixed points on the phase plain, we numerically solved the equation 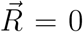 starting from a few different initial conditions using Matlab fsolve function. To find the stable and unstable manifolds of the saddle, we followed the path along −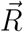 and 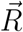, respectively, starting the trajectory near the saddle point.

## Acknowledgements

This work was supported by the Swartz Foundation (M.G.), National Institutes of Health (NIH) grant R01 EB026949 (T.A.E. and M.G.), NIH grant RF1DA055666 (M.G. and T.A.E.), NIH grant S10OD028632-01 (M.G. and T.A.E.), Alfred P. Sloan Foundation Research Fellowship (T.A.E.), NIH grant K99/R00 NS092972 (C.C.), NIH grant NS121409 (C.C.), NIH grant NS122969 (C.C.), the Brain and Behavior Research Foundation (C.C.), and the Whitehall Foundation (C.C.). We thank C. Aghamohammadi for sharing the methods on point process variability and J. Roach for sharing the code for the analysis of the mean-field network model.

## Author contributions

M.G. and T.A.E. designed the research and developed the computational analysis framework. C.C. and K.S. designed the experiments. M.G. developed the code, performed computer simulations, and analysed neural recordings data. C.C. performed the experiments, spike sorting, and data curation. M.G., T.A.E and C.C. wrote the paper.

## Data availability

The synthetic data used in this study can be reproduced using the source code. Neural recordings data will be made publicly available on Figshare upon publication.

## Code availability

The source code to reproduce the results of this study will be made available on GitHub upon publication.

## References

1. Cunningham, J. P. & Yu, B. M. Dimensionality reduction for large-scale neural recordings. Nat. Neurosci. 17, 1500–1509 (2014).

2. Chang, L. & Tsao, D. Y. The code for facial identity in the primate brain. Cell 169, 1013–1028.e14 (2017).

3. Gallego, J. A., Perich, M. G., Miller, L. E. & Solla, S. A. Neural manifolds for the control of movement. Neuron 94, 978–984 (2017).

4. Chaudhuri, R., Gerçek, B., Pandey, B., Peyrache, A. & Fiete, I. The intrinsic attractor manifold and population dynamics of a canonical cognitive circuit across waking and sleep. Nat. Neurosci. 22, 1512–1520 (2019).

5. Gardner, R. J. et al. Toroidal topology of population activity in grid cells. Nature 602, 123–128 (2022).

6. Jazayeri, M. & Ostojic, S. Interpreting neural computations by examining intrinsic and embedding dimensionality of neural activity. Curr. Opin. Neurobiol. 70, 113–120 (2021).

7. Langdon, C., Genkin, M. & Engel, T. A. A unifying perspective on neural manifolds and circuits for cognition. Nat. Rev. Neurosci. 24, 363–377 (2023).

8. Kriegeskorte, N. & Wei, X.-X. Neural tuning and representational geometry. Nat. Rev. Neurosci. 22, 703–718 (2021).

9. Nieh, E. H. et al. Geometry of abstract learned knowledge in the hippocampus. Nature 595, 80–84 (2021).

10. Bollimunta, A., Totten, D. & Ditterich, J. Neural dynamics of choice: Single-trial analysis of decision-related activity in parietal cortex. J. Neurosci. 32, 12684–12701 (2012).

11. Latimer, K. W., Yates, J. L., Meister, M. L. R., Huk, A. C. & Pillow, J. W. Single-trial spike trains in parietal cortex reveal discrete steps during decision-making. Science 349, 184–187 (2015).

12. Cohen, M. R. & Maunsell, J. H. R. A neuronal population measure of attention predicts behavioral performance on individual trials. J. Neurosci. 30, 15241–15253 (2010).

13. Engel, T. A. et al. Selective modulation of cortical state during spatial attention. Science 354, 1140–1144 (2016).

14. Denfield, G. H., Ecker, A. S., Shinn, T. J., Bethge, M. & Tolias, A. S. Attentional fluctuations induce shared variability in macaque primary visual cortex. Nat. Commun. 9, 334–14 (2018).

15. Chandrasekaran, C., Peixoto, D., Newsome, W. T. & Shenoy, K. V. Laminar differences in decision-related neural activity in dorsal premotor cortex. Nat. Commun. 8, 996 (2017).

16. Mante, V., Sussillo, D., Shenoy, K. V. & Newsome, W. T. Context-dependent computation by recurrent dynamics in prefrontal cortex. Nature 503, 78–84 (2013).

17. Murray, J. D. et al. Stable population coding for working memory coexists with heterogeneous neural dynamics in prefrontal cortex. Proc. Natl. Acad. Sci. U.S.A 114, 394–399 (2017).

18. Chaisangmongkon, W., Swaminathan, S. K., Freedman, D. J. & Wang, X.-J. Computing by robust transience: How the fronto-parietal network performs sequential, category-based decisions. Neuron 93, 1504–1517.e4 (2017).

19. Cavanagh, S. E., Towers, J. P., Wallis, J. D., Hunt, L. T. & Kennerley, S. W. Reconciling persistent and dynamic hypotheses of working memory coding in prefrontal cortex. Nat. Commun. 9, 3498 (2018).

20. Chandrasekaran, C. et al. Brittleness in model selection analysis of single neuron firing rates. bioRxiv preprint at https://www.biorxiv.org/content/10.1101/430710v1 (2018).

21. Zoltowski, D. M., Latimer, K. W., Yates, J. L., Huk, A. C. & Pillow, J. W. Discrete stepping and nonlinear ramping dynamics underlie spiking responses of lip neurons during decision-making. Neuron 102, 1249–1258.e10 (2019).

22. Steinemann, N. A. et al. Direct observation of the neural computations underlying a single decision. bioRxiv preprint at https://www.biorxiv.org/content/10.1101/2022.05.02.490321v3 (2023).

23. Genkin, M. & Engel, T. A. Moving beyond generalization to accurate interpretation of flexible models. Nat. Mach. Intell. 2, 674–683 (2020).

24. Genkin, M., Hughes, O. & Engel, T. A. Learning non-stationary Langevin dynamics from stochastic observations of latent trajectories. Nat. Commun. 12, 5986 (2021).

25. Zoltowski, D. M., Pillow, J. W. & Linderman, S. W. Unifying and generalizing models of neural dynamics during decision-making. arXiv preprint at https://arxiv.org/abs/2001.04571 (2020).

26. Pandarinath, C. et al. Inferring single-trial neural population dynamics using sequential auto-encoders. Nat. Methods 15, 805–815 (2018).

27. Gold, J. I. & Shadlen, M. N. The neural basis of decision making. Annu. Rev. Neurosci. 30, 535–574 (2007).

28. Peixoto, D. et al. Decoding and perturbing decision states in real time. Nature 80, 791–21 (2021).

29. Aghamohammadi, C. & Engel, T. A. Unbiased estimation of firing-rate variance from spikes to reveal decision computations. 48th Annual Meeting of the Society for Neuroscience (2019). A bioRxiv preprint for this work will be submitted shortly.

30. Wang, X.-J. Probabilistic decision making by slow reverberation in cortical circuits. Neuron 36, 955–968 (2002).

31. Wong, K.-F. & Wang, X.-J. A recurrent network mechanism of time integration in perceptual decisions. J. Neurosci. 26, 1314–1328 (2006).

32. Song, H. F., Yang, G. R. & Wang, X.-J. Training excitatory-inhibitory recurrent neural networks for cognitive tasks: A simple and flexible framework. PLoS Comput. Biol. 12, e1004792–30 (2016).

33. Mastrogiuseppe, F. & Ostojic, S. Linking connectivity, dynamics, and computations in low-rank recurrent neural networks. Neuron 99, 609–623.e29 (2018).

34. Inagaki, H. K., Fontolan, L., Romani, S. & Svoboda, K. Discrete attractor dynamics underlies persistent activity in the frontal cortex. Nature 566, 212–217 (2019).

35. Finkelstein, A. et al. Attractor dynamics gate cortical information flow during decision-making. Nat. Neurosci. 24, 843–850 (2021).

36. Roach, J. P., Churchland, A. K. & Engel, T. A. Choice selective inhibition drives stability and competition in decision circuits. Nat. Commun. 14, 147 (2023).

37. Langdon, C. & Engel, T. A. Latent circuit inference from heterogeneous neural responses during cognitive tasks. bioRxiv preprint at https://www.biorxiv.org/content/10.1101/2022.01.23.477431v1 (2022).

38. Yang, T. & Shadlen, M. N. Probabilistic reasoning by neurons. Nature 447, 1075–1080 (2007).

39. Hanks, T. D. et al. Distinct relationships of parietal and prefrontal cortices to evidence accumulation. Nature 520, 220–223 (2015).

40. Kingma, D. P. & Ba, J. Adam: A method for stochastic optimization. arXiv preprint at https://arxiv.org/abs/1412.6980 (2014).

41. Okuta, R., Unno, Y., Nishino, D., Hido, S. & Loomis, C. CuPy: A NumPy-compatible library for NVIDIA GPU calculations. In Proceedings of Workshop on Machine Learning Systems (LearningSys) in The Thirty-first Annual Conference on Neural Information Processing Systems (NIPS) (2017).

42. Hastie, T., Tibshirani, R. & Friedman, J. The Elements of Statistical Learning (Springer Science & Business Media, 2013).

43. Belkin, M., Hsu, D., Ma, S. & Mandal, S. Reconciling modern machine-learning practice and the classical bias–variance trade-off. Proc. Natl. Acad. Sci. U.S.A 116, 15849–15854 (2019).

44. Haas, K. R., Yang, H. & Chu, J.-W. Trajectory entropy of continuous stochastic processes at equilibrium. J. Phys. Chem. Lett. 5, 999–1003 (2014).

45. Brown, E. N., Barbieri, R., Ventura, V., Kass, R. E. & Frank, L. M. The time-rescaling theorem and its application to neural spike train data analysis. Neural Comput. 14, 325–346 (2002).

46. Cox, D. R. Renewal Theory (Springer, 1967).

47. Cox, D. R. & Isham, V. Point Processes (CRC Press, 1980).

48. Cox, D. R. & Lewis, P. A. The Statistical Analysis of Series of Events (Springer, 1966).

49. Maimon, G. & Assad, J. A. Beyond Poisson: Increased spike-time regularity across primate parietal cortex. Neuron 62, 426–440 (2009).

50. Bishop, C. M. Pattern Recognition and Machine Learning (Springer, New York, 2007).

51. Stimberg, M., Brette, R. & Goodman, D. F. Brian 2, an intuitive and efficient neural simulator. eLife 8, e47314 (2019).

