## Supplementary Information for "The dynamics and geometry of choice in premotor cortex"

July 22, 2023

**3 Supplementary References** **20**

### 1 Supplementary Figures

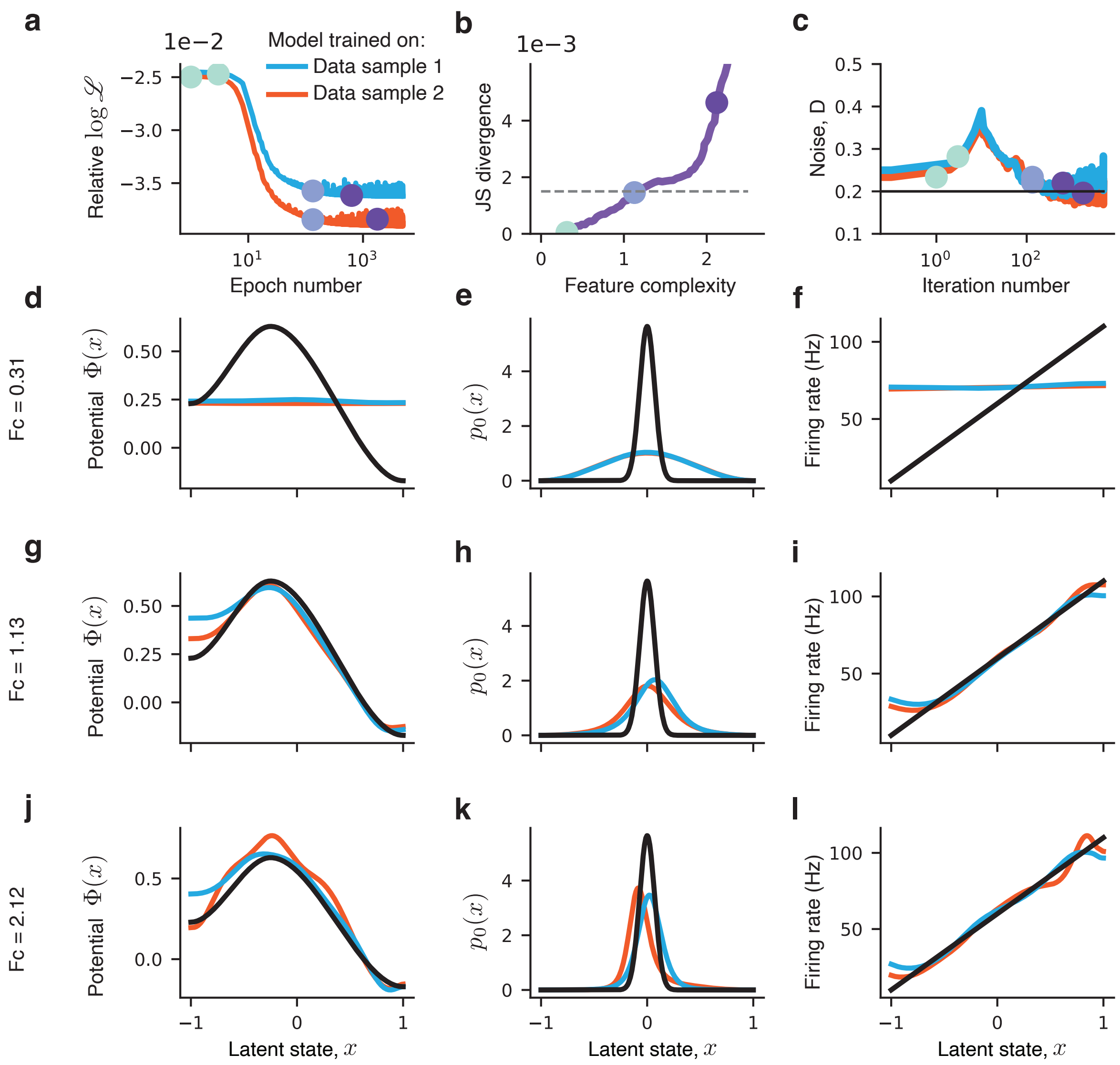

**Supplementary Figure 1. Model fitting and model selection on synthetic data.** We generated 400 trials of spike data for a single neuron from a ground-truth model with a single-barrier potential, narrow zero-centered $p_0(x)$ distribution and linear firing rate function. We fit the model independently on two halves of the data $\mathcal{D}_1$ and $\mathcal{D}_2$ (200 trials each) and compare features between the models inferred from each data halve to select the optimal model. **a**, Relative log-likelihood ($[\log \mathscr{L}_{\mathrm{gt}} - \log \mathscr{L}] \,/\, \log \mathscr{L}_{\mathrm{gt}}$, where $\mathscr{L}_{\mathrm{gt}}$ is the likelihood of the ground truth model) decreases with the epoch number indicating an improving fit on the training data. **b**, Jensen-Shannon (JS, y-axis) divergence between the models with the same feature complexity (FC, x-axis) inferred from two data halves. Colored dots indicate three levels of FC that illustrate underfitting (mint), optimal (blue), and overfitting (purple) regimes. The optimal model is selected at FC where JS divergence exceeds a fixed threshold $\mathrm{JS}_{\mathrm{thres}} = 0.0015$. The JS threshold was the same for all results in this work. **c**, Noise magnitude $D$ for each training epoch for models fitted on $\mathcal{D}_1$ and $\mathcal{D}_2$. Black line indicates the ground-truth value of $D$. **d**, The ground-truth potential (black) and the potentials of the models fitted on $\mathcal{D}_1$ and $\mathcal{D}_2$ (colored lines) for low feature complexity (mint dot in b). The inferred potentials overlap but miss some of the ground-truth features, which is the underfitting regime. **e**, Same as d for the initial state distribution $p_0(x)$. **f**, Same as d for the tuning function $f(x)$. **g-i** Same as d-f for the optimal feature complexity (blue dot in b). The models inferred from two data halves overlap and match the ground-truth model. **j-l** Same as d-f for high feature complexity (purple dot in b). The models inferred from two data halves diverge and disagree with the ground-truth, which is the overfitting regime.

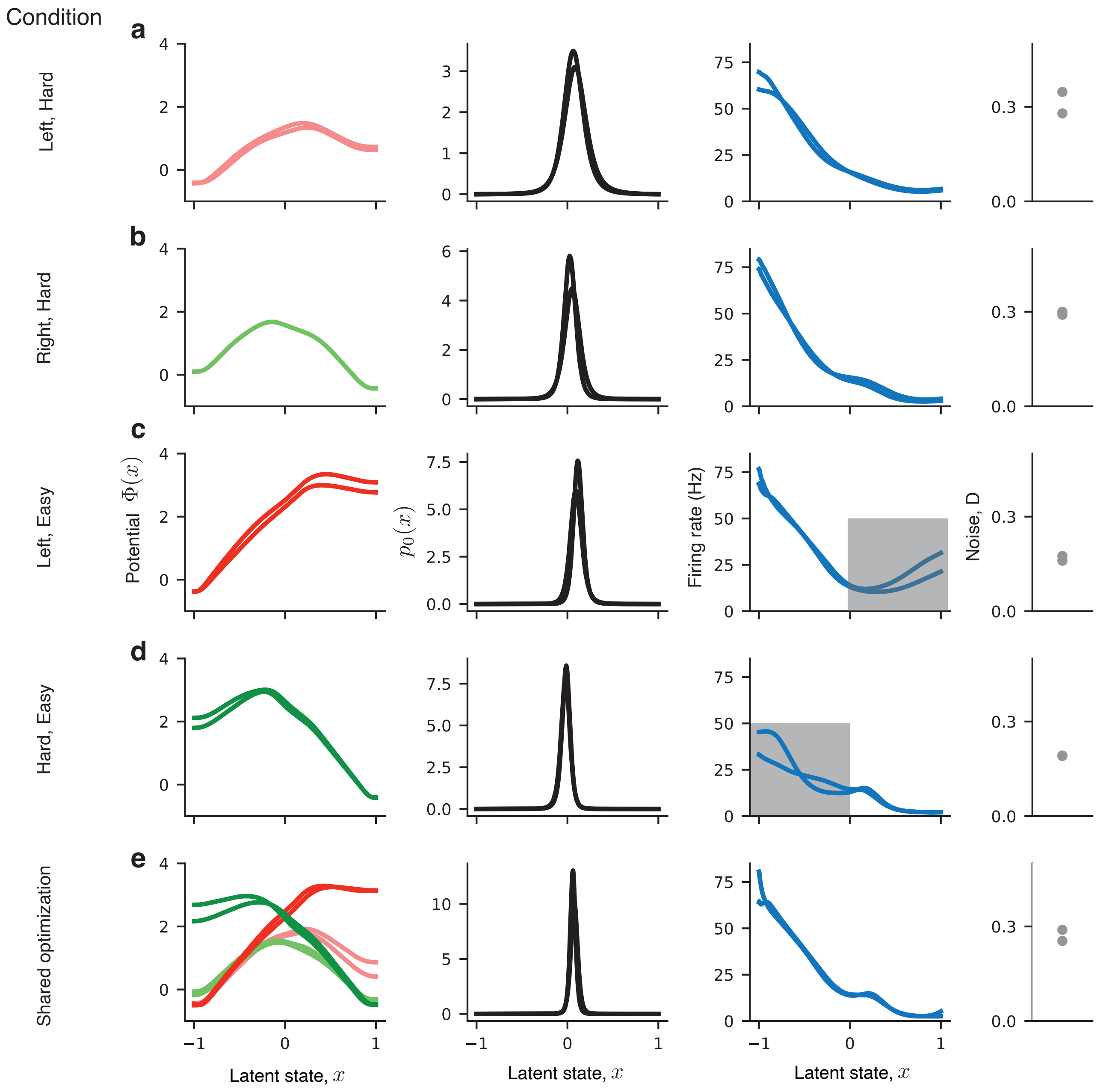

**Supplementary Figure 2. Independent and shared optimizations across four stimulus conditions for an example PMd neuron. a**, Fitted potential functions (left), $p_0(x)$ distributions (middle left), tuning functions (middle right) and noise magnitudes (right) for a left-hard condition obtained from two data halves $\mathcal{D}_1$ and $\mathcal{D}_2$. This model was selected using our model selection method based on feature consistency. **b**, Same as a for the right-hard condition. The slope of the inferred potential points towards the opposite boundary than in the left-hard condition, while the inferred tuning function is the same. **c**, Same as a for the left-easy condition. The inferred tuning function is the same as in the hard conditions in the left part of the domain where the dynamics evolve towards the correct choice, but the tuning function is inferred less accurately in the right side of the domain (grey highlight) due to very small number of rightward choices (error trials) in the left-easy stimulus condition. **d**, Same as c for the right-easy condition. **e**, Shared optimization across all four stimulus conditions, in which $p_0(x)$, tuning functions, and noise magnitude are restricted to be the same and only the potential $\Phi(x)$ can vary across stimulus conditions. The shared optimization enables more accurate inference of tuning functions, because it learns a single tuning function across four conditions so that the number of leftward and rightward choices are approximately balanced in the data, and the dynamics equally explore both sides of the decision manifold.

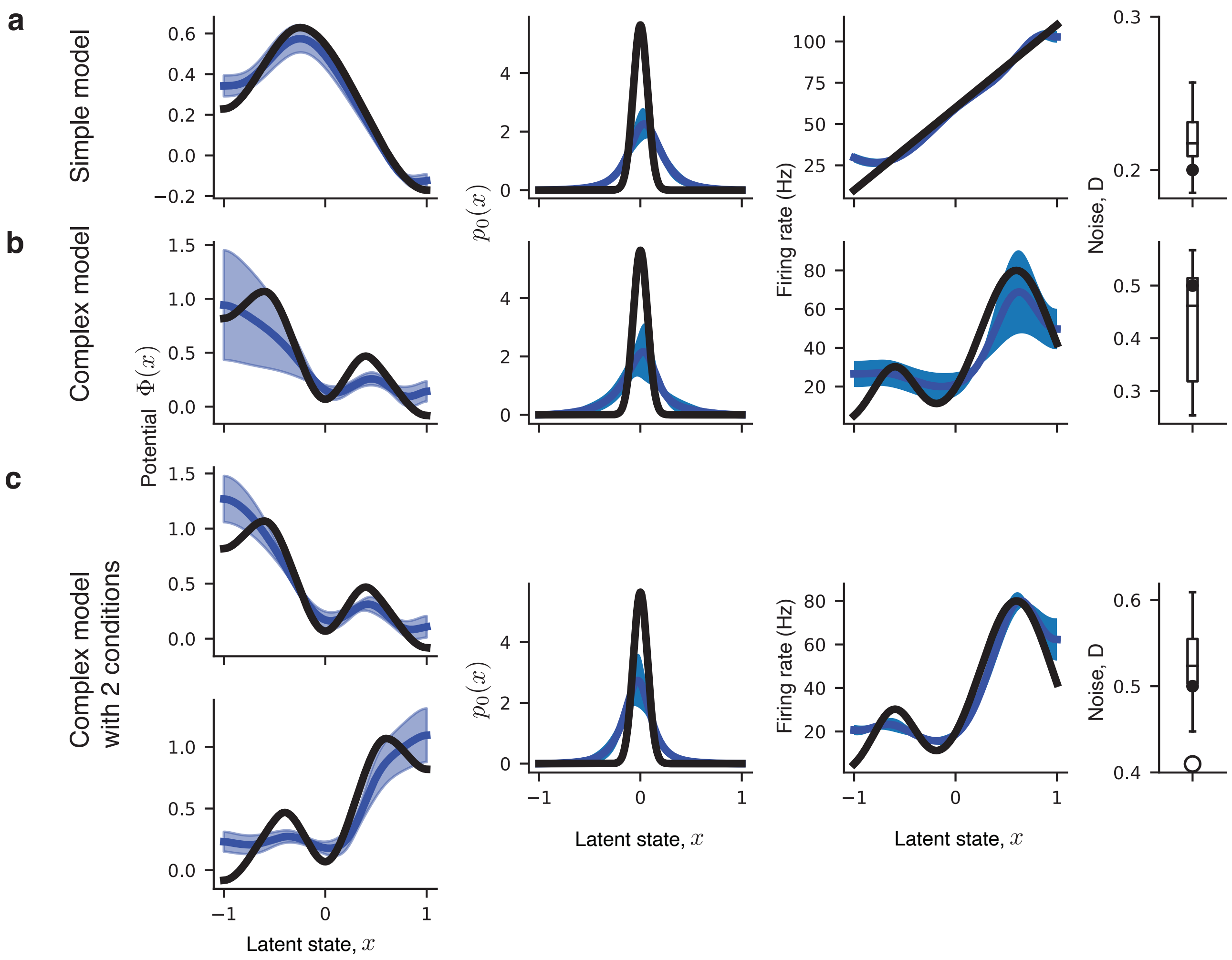

**Supplementary Figure 3. Inference of dynamics with different complexity from synthetic single-neuron data.** We generate spike data for a single neuron using our model with varying complexity of the potential and tuning function, with the amount of data (800 trials total) similar to our PMd recordings. We perform the model fitting and model selection to test how the inference accuracy depends on the complexity of the ground-truth dynamics. **a**, The inferred potential (left), $p_0(x)$ distribution (middle left), tuning function (middle right) and noise magnitude (right) for a model with a single-barrier potential and a linear tuning function. The inferred model (blue) tightly overlaps with the ground-truth (black). **b**, Same as a for a model that has a more complex potential with two barriers and a non-linear tuning function. The inferred model matches the ground truth, but the estimation uncertainty is higher and the inference is less accurate in the regions that are poorly explored by the dynamics in the data (the region with high potential near the left boundary). The inference accuracy improves when fitting data from a population of neurons (Supplementary Fig. 4). **c**, Shared optimization across two stimulus conditions for the same model as in b. In two conditions, the potentials are mirror images of each other, and $p_0(x)$, $D$ and tuning function are the same. In the shared optimization, we restrict $p_0(x)$, $D$ and the tuning function to be the same across conditions. The shared optimization leads to a lower estimation uncertainty and more accurate inference (cf. to b). The inference accuracy further improves when fitting data from a population of neurons using shared optimization (Supplementary Fig. 5). In all panels, error bars are s.t.d. across 10 bootstrap samples.

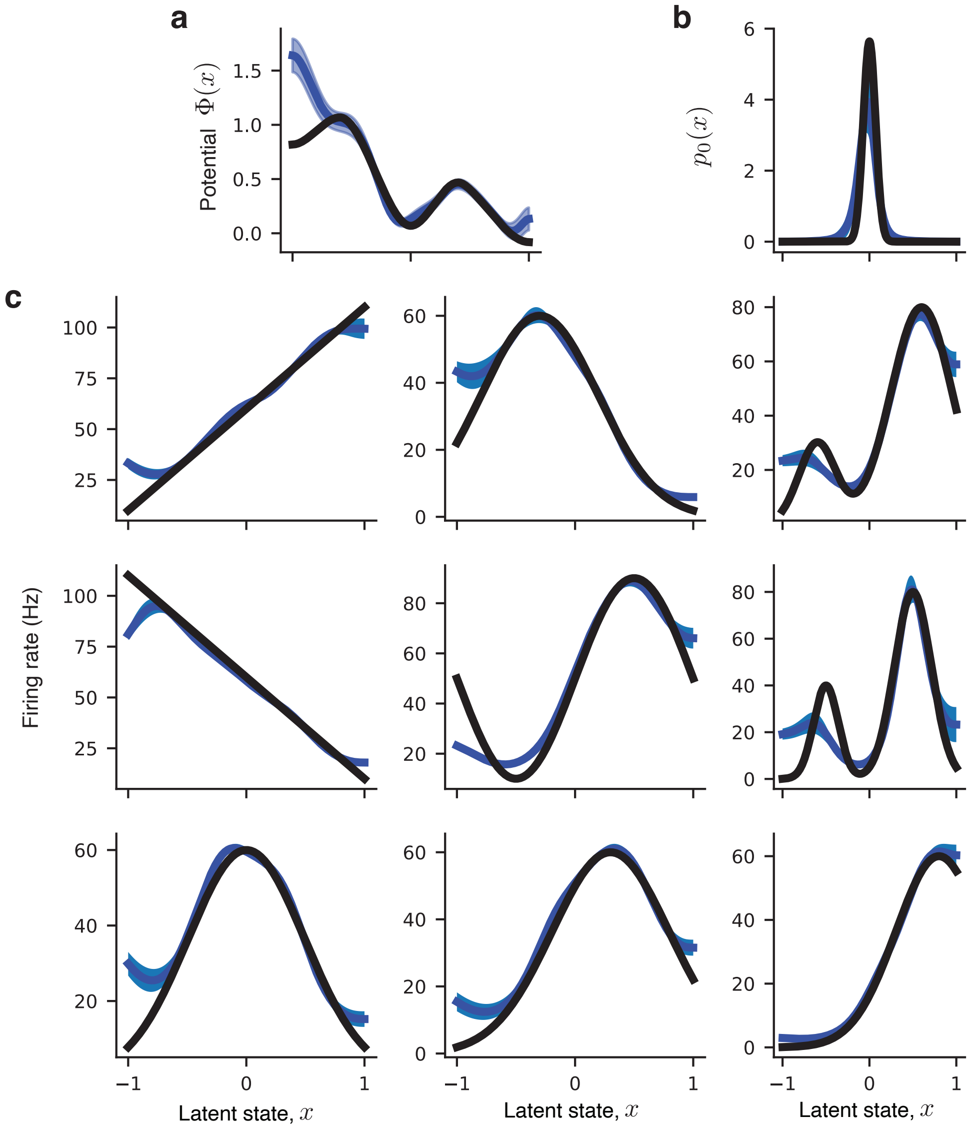

**Supplementary Figure 4. Inference of complex dynamics from synthetic population data.** We generate spike data for a population of 9 neurons using our model with a complex two-barrier potential (same is in Supplementary Fig. 3b) and heterogeneous non-linear tuning functions, with the amount of data (800 trials total) similar to our PMd recordings. We perform the model fitting and model selection to test the inference accuracy. **a**, The inferred potential (blue) tightly overlaps with the ground truth (black). The inference is inaccurate near the left boundary, since this region with the high potential is poorly explored by the dynamics in the data. The inference accuracy further improves for the shared optimization across multiple stimulus conditions (Supplementary Fig. 5). **b**, The inferred $p_0(x)$ distribution (blue) tightly overlaps with the ground truth (black). **c**, The inferred tuning functions (blue) tightly overlap with the ground truth (black). The inference of tuning functions is more accurate compared to the single-neuron models (cf. Supplementary Fig. 3), because spikes of multiple neurons in the population contribute to a more precise estimation of the latent state, which in turn enables a more accurate inference of the tuning function of each neuron. In all panels, error bars are s.t.d. across 10 bootstrap samples.

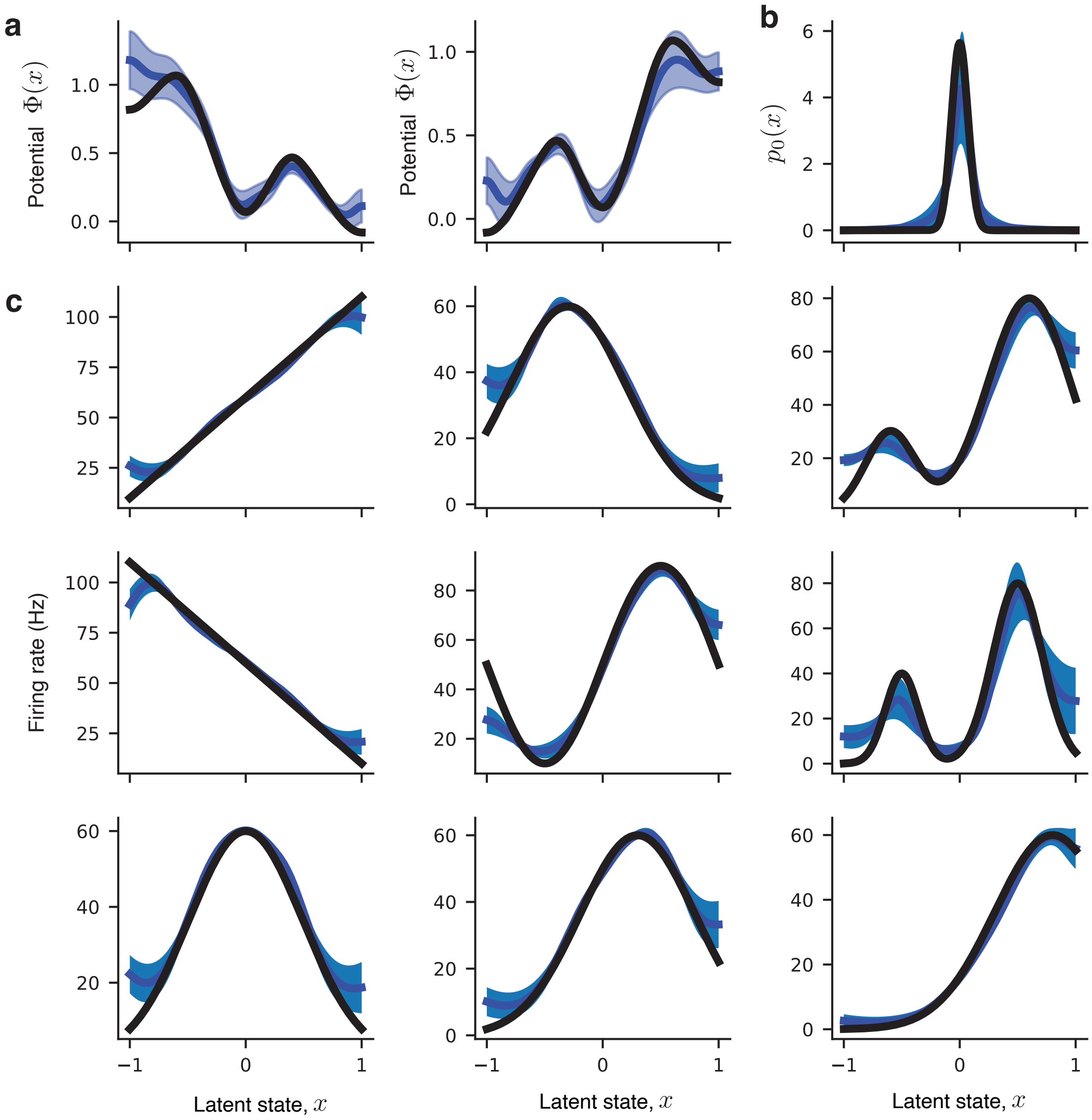

**Supplementary Figure 5. Shared inference of complex dynamics from synthetic population data.** We generate spike data for a population of 9 neurons using the same model as in Supplementary Fig. 4 for two stimulus conditions. In two conditions, the potentials are mirror images of each other, and $p_0(x)$, $D$ and tuning function are the same. In the shared optimization, we restrict $p_0(x)$, $D$ and the tuning functions to be the same across conditions. We perform the model fitting and model selection to test the inference accuracy. **a**, The inferred potentials (blue) tightly overlap with the ground truth (black) in both conditions. The inference is accurate even in the regions with high potential in each condition, because the tuning functions inferred from one condition enable more accurate inference of dynamics from limited data in the other condition. **b**, The inferred $p_0(x)$ distribution (blue) tightly overlaps with the ground truth (black). **c**, The inferred tuning functions (blue) tightly overlap with the ground truth (black). The inference is accurate on both left and right sides of the latent space, since the regions not explored by the dynamics in one condition are explored in another condition, which enables the accurate inference of tuning functions in the entire domain.

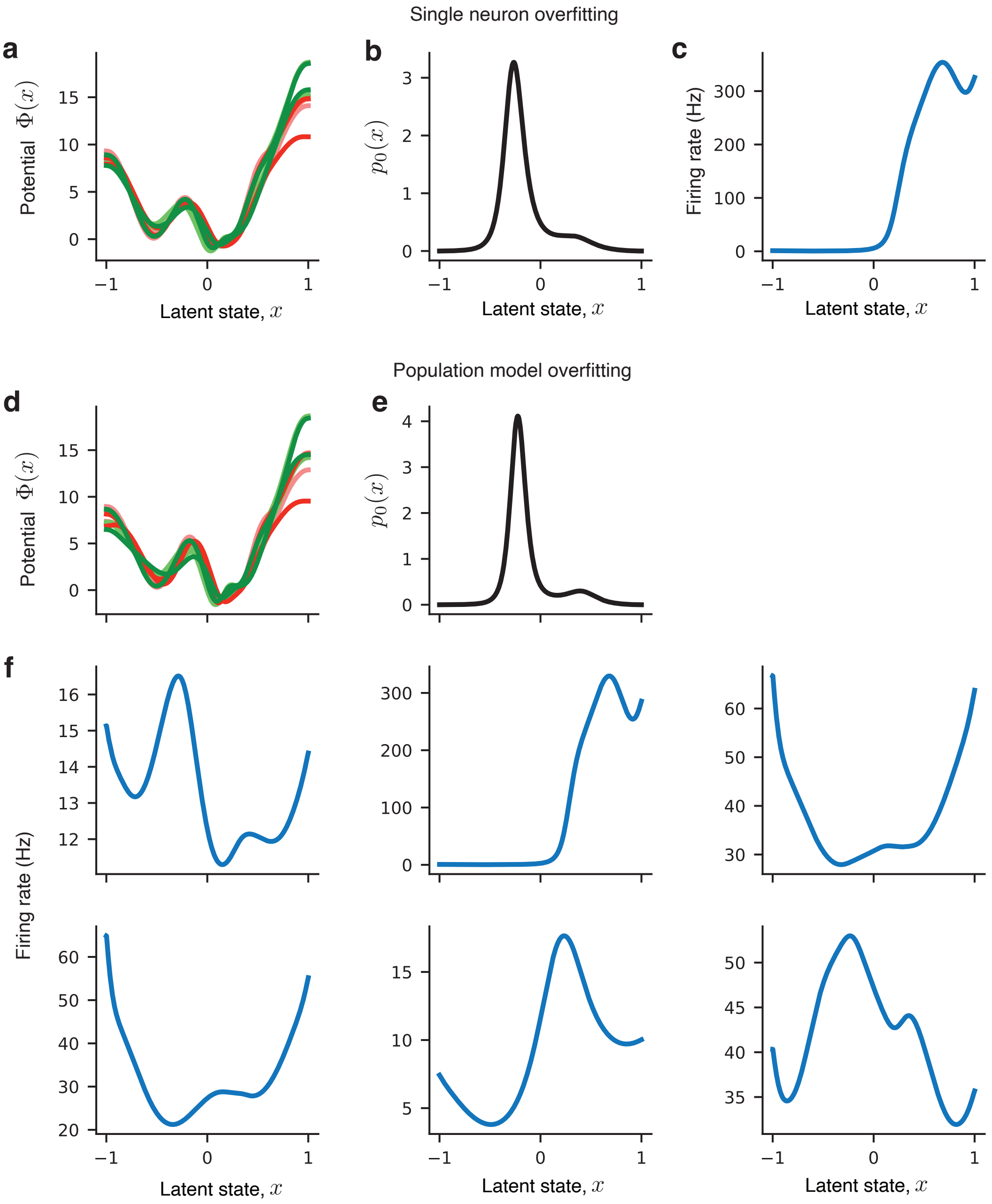

**Supplementary Figure 6. Overfitting on PMd data.** When fitting our model to PMd data and performing model selection, we observed three instances of overfitting: 1 single neuron and 1 population from monkey O, and 1 population from monkey T. **a**, The model showing overfitting for the single neuron from monkey O. The inferred potentials are the same across four stimulus conditions and show deep wells compensated by a disproportionally high noise magnitude ($D \sim 3-5$, compared to $D \sim 0.2-0.6$ in regular fits). This model produces severely underestimated reaction times (reaction time ∼10 ms in the model, compared to ∼500 ms in the data) and does not predict monkey's choice. **b**, The inferred $p_0(x)$ shared across conditions for the model in a. **c**, The inferred tuning function shared across conditions for the model in a shows unrealistically high firing rates up to several hundreds of Hz. **d**, The model showing overfitting for the population from monkey O. The inferred potentials are the same across four stimulus conditions and show deep wells compensated by a disproportionally high noise magnitude ($D \sim 4$). This model produces severely underestimated reaction times and does not predict monkey's choice. The overfitted model for the population from monkey T had similar features (not shown here). **e**, The inferred $p_0(x)$ shared across conditions for the model in d. **f**, The inferred tuning functions shared across conditions for the model in d. The single neuron shown in a-c is part of this population (upper row, middle), and its tuning function shows unrealistically high firing rates in the population model as well.

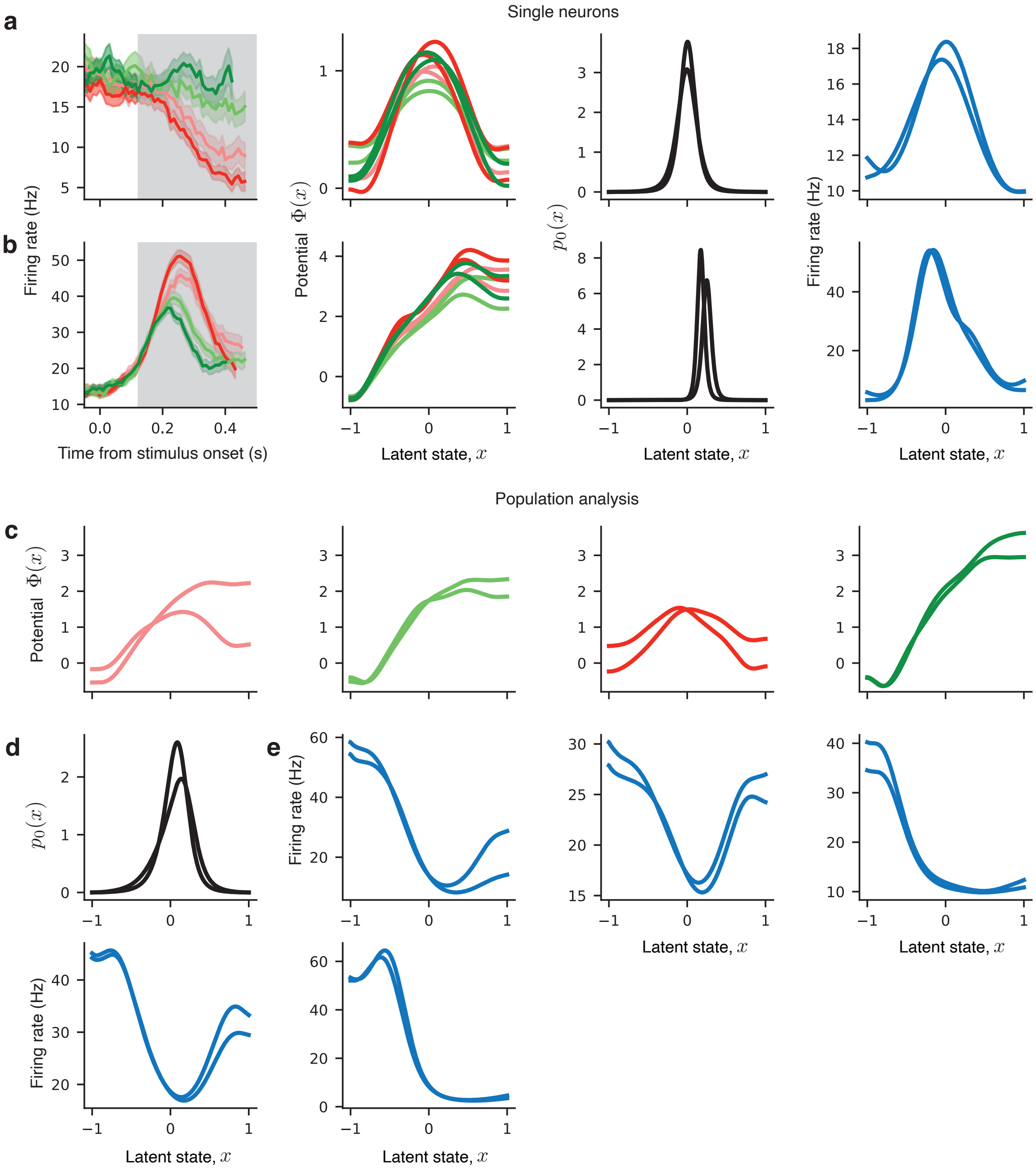

**Supplementary Figure 7. Underfitting on PMd data.** We observed two types of underfitting: no decision signal (a,b) and disagreement between data splits (c-e), both of which may arise when a model cannot detect a weak decision signal and mainly fits the condition-independent trend in neural activity. **a**, A model showing underfitting for an example single neuron. Left to right: trial-average firing rates sorted by the chosen side, the inferred potentials for four stimulus conditions, $p_0(x)$ and the tuning function shared across conditions. The potentials have a similar symmetric shape across all conditions, such that the left and right choices are equally probable, and the tuning function is also symmetric. This model captures only the overall ramping trend but not the decision signal, which likely results from weak choice selectivity of this neuron. **b**, Same as a for another example neuron. In all conditions, the potentials point to the left boundary, predicting more left choices in all conditions. This model captures the speed of the dynamics (steeper slope for easy conditions) but not the decision signal. **c**, A model showing underfitting for an example population of 6 neurons. The potentials inferred from two data halves in four stimulus conditions. In easy-left condition (third panel from the left), the potentials disagree between the two data splits pointing to the opposite boundaries, which leads to an early crossing of JS divergence threshold and the selection of a model with low feature complexity before all consistent features have been discovered. **d**, The inferred $p_0(x)$ shared across conditions from the model in c. **e**, The inferred tuning functions shared across conditions from the model in c.

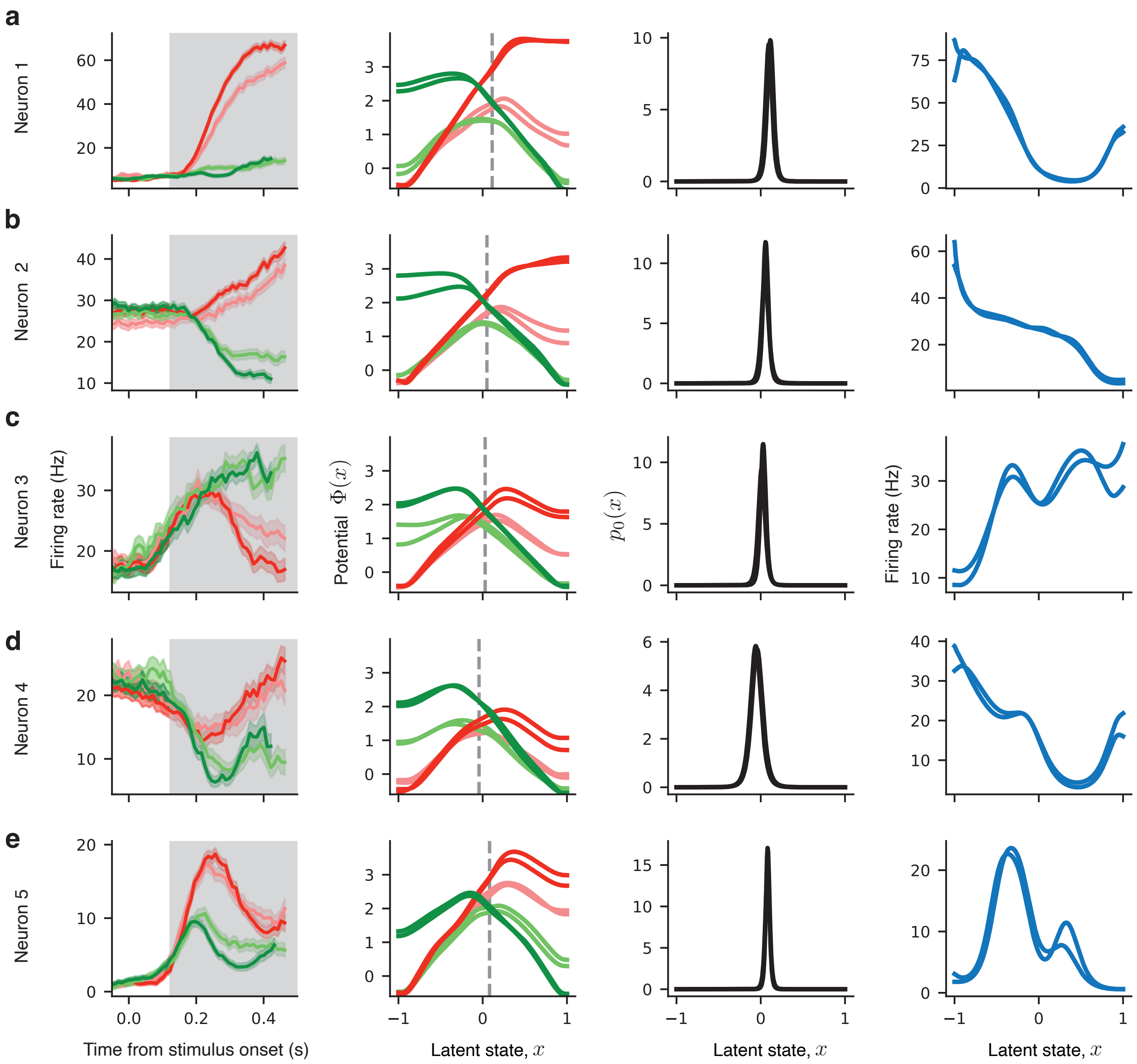

**Supplementary Figure 8. The inferred models with a single-barrier potential for additional example single neurons in PMd. a**, Trial-average firing rates sorted by the chosen side and stimulus difficulty (left), the inferred potentials for four stimulus conditions (middle left), $p_0(x)$ distribution shared across conditions (middle right), and tuning functions shared across conditions (right) for a single PMd neuron. **b-e**, Same as a for four other example neurons. Despite heterogeneous profiles of the trial-average firing rates, all models show the same dynamics described by a single-barrier potential and narrow zero-centered $p_0(x)$ distribution. The response heterogeneity results from diverse tuning functions to the latent variable $x$.

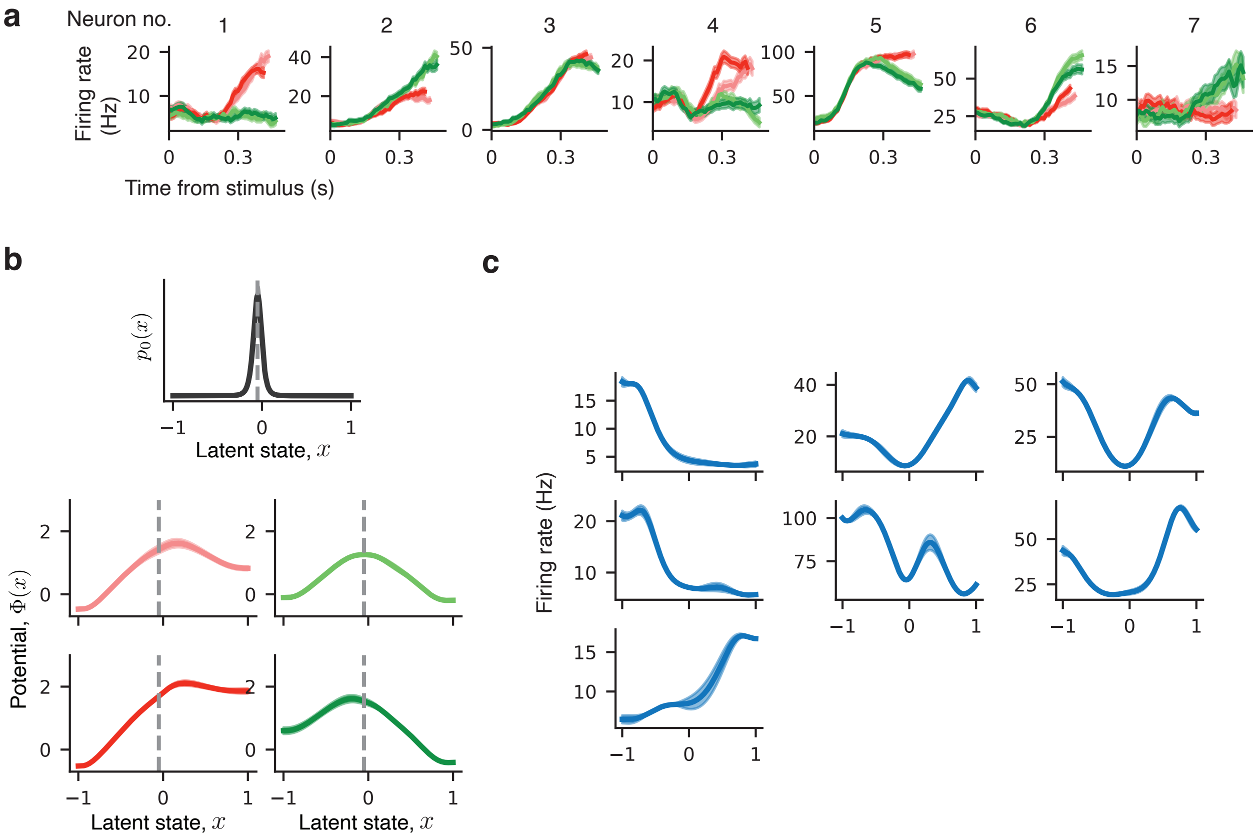

**Supplementary Figure 9. The inferred model with a single-barrier potential for additional example population of PMd neurons. a**, Trial-average firing rates sorted by the chosen side and stimulus difficulty for a population of 7 neurons recorded simultaneously from monkey O. **b**, The inferred potentials for four stimulus conditions (middle and lower panels) and $p_0(x)$ distribution shared across conditions (upper panel) for the population in a. **c**, The inferred tuning functions shared across conditions for the population in a.

#### 2 Supplementary Materials and Methods

##### 2.1 Analytical calculation and numerical evaluation of the likelihood

The likelihood is calculated by marginalizing the joint probability of simultaneously observing a latent trajectory $\mathcal{X}(t)$ and spike data $Y(t)$ from a model $\theta$:

$$\mathscr{L}\left[Y(t)|\theta\right] = \int \mathscr{D}\mathcal{X}(t)\; P(\mathcal{X}(t), Y(t)|\theta). \tag{1}$$

We first integrate $P(\mathcal{X}(t), Y(t)|\theta)$ over the distribution of all latent paths in between the observed spikes to obtain the joint probability $P(X(t), Y(t)|\theta)$ where $X(t) = \{x_{t_0}, x_{t_1}, \dots, x_{t_N}, x_{t_E}\}$ is a discretized trajectory which consists of the initial state $x_{t_0}$, the final state at the trial end $x_{t_E}$, and all states $x_{t_1}, ...x_{t_N}$ at the times of spike observations from all neurons. The likelihood is obtained by marginalizing $P(X(t), Y(t)|\theta)$ over the discretized trajectory:

$$\mathscr{L}\left[Y(t)|\theta\right] = \int_{x_{t_0}} \int_{x_{t_1}} \cdots \int_{x_{t_N}} \int_{x_{t_E}} dx_{t_0} \dots dx_{t_E} P(X(t), Y(t)|\theta). \tag{2}$$

Using the Markov property of the latent Langevin dynamics Eq. (1) and conditional independence of spike observations, the joint probability density $P(X(t), Y(t))$ can be factorized [1]:

$$P(X(t), Y(t)) = p(x_{t_0}) \left( \prod_{i=1}^{N} p(y_{t_i}|x_{t_i}) p(x_{t_i}|x_{t_{i-1}}) \right) p(x_{t_E}|x_{t_N}) p(A|x_{t_E}). \tag{3}$$

Here $p(y_{t_i}|x_{t_i})dt$ is the probability of observing a spike from neuron $k_i$ within a small $dt$ of time $t_i$ given the latent state $x_{t_i}$, hence $p(y_{t_i}|x_{t_i}) = f_{k_i}(x_{t_i})$ by the definition of the instantaneous Poisson firing rate, where $k_i$ is the index of the neuron that emitted a spike at time $t_i$. $p(x_{t_0})$ is the probability density of the initial latent state. $p(x_{t_i}|x_{t_{i-1}})$ is the transition probability density from $x_{t_{i-1}}$ to $x_{t_i}$ during the time interval between the adjacent spike observations from all neurons, which accounts for the absence of spikes during this time interval. This transition probability is marginalized over all intermediate latent paths connecting $x_{t_{i-1}}$ at time $t_{i-1}$ and $x_{t_i}$ at time $t_i$. Finally, the term $p(A|x_{t_E})$ is the absorption operator, which ensures that only trajectories terminating at one of the domain boundaries at time $t_E$ contribute to the likelihood [1]. $p(x_{t_i}|x_{t_{i-1}})$ is a solution of the modified Fokker-Planck equation [1]:

$$\frac{\partial p(x,t)}{\partial t} = \left( -D\frac{\partial}{\partial x} F(x) + D\frac{\partial^2}{\partial x^2} - \sum_{k=1}^{M} f_k(x) \right) p(x,t) \equiv -\hat{\boldsymbol{\mathcal{H}}} p(x,t), \tag{4}$$

where the term $\sum_{k=1}^{M} f_k(x)$ is the total firing rate of all neurons that accounts for the absence of spikes during the interspike intervals. The absorbtion operator is $\boldsymbol{A} = \hat{\boldsymbol{\mathcal{H}}}_0$, where $\hat{\boldsymbol{\mathcal{H}}}_0$ is the Fokker-Planck operator [1]:

$$\hat{\boldsymbol{\mathcal{H}}}_0 = D\frac{\partial}{\partial x} F(x) - D\frac{\partial^2}{\partial x^2}. \tag{5}$$

To numerically calculate the likelihood using Eq. (2), we need to solve Eq. (4) for each interspike interval. To solve Eq. (4), we introduce a Hermitian operator $\boldsymbol{\mathcal{H}} = \exp(\Phi(x)/2)\hat{\boldsymbol{\mathcal{H}}}\exp(-\Phi(x)/2)$ that propagates forward in time the scaled probability density $\rho(x,t) = p(x,t)\exp(\Phi(x)/2)$:

$$\frac{\partial \rho(x,t)}{\partial t} = -\boldsymbol{\mathcal{H}}\rho(x,t). \tag{6}$$

Eq. (6) is a linear equation with a symmetric differential operator that allows for an efficient numerical solution. The operator $\boldsymbol{\mathcal{H}}$ can be decomposed into a sum of two operator: $\boldsymbol{\mathcal{H}} = \boldsymbol{\mathcal{H}}_0 + \boldsymbol{\mathcal{H}}_I$, where $\boldsymbol{\mathcal{H}}_0$ accounts for drift and diffusion in the latent space, and $\boldsymbol{\mathcal{H}}_I$ accounts for the probability decay during interspike intervals:

$$\begin{aligned}\boldsymbol{\mathcal{H}}_0 &= -e^{\Phi(x)/2}\frac{\partial}{\partial x}De^{-\Phi(x)}\frac{\partial}{\partial x}e^{\Phi(x)/2},\\ \boldsymbol{\mathcal{H}}_I &= \sum_{k=1}^{M} f_k(x).\end{aligned} \tag{7}$$

First, we find the eigenvalues and eigenvectors of the operator $\boldsymbol{\mathcal{H}}_0$:

$$\boldsymbol{\mathcal{H}}_0\Psi_0(x) = \lambda\Psi_0(x). \tag{8}$$

To this end, we introduce scaled eigenfunctions $\phi_0(x) = \exp(\Phi(x)/2)\Psi_0(x)$, which are the solution of the following scaled eigenvalue problem:

$$-\frac{\partial}{\partial x}De^{-\Phi(x)}\frac{\partial}{\partial x}\phi_0(x) = \lambda_0 e^{-\Phi(x)}\phi_0(x). \tag{9}$$

We solve the problem Eq. (9) numerically using the spectral elements method (SEM, see Supplementary Information in Refs. [1, 2] for details). We obtain the set of eigenvalues $\lambda_{0,i}$ and the eigenvectors $\phi_{0,i}(x_k)$, where $i = 1, 2, ..., N_v$ indexes the eigenvalues and eigenvectors, $N_v$ is the number of retained eigenvalues, and $k$ indexes grid points in the discretized domain. For all model fits, we set $N_v = N - 2 = 447$, where $N = 449$ is the size of the SEM grid. The problem Eq. (9) is a generalized eigenvalue problem with a non-trivial right-hand side function $\exp(-\Phi(x))$. With the SEM discretization, the mass matrix $\boldsymbol{M}$ for Eq. (9) is a product of $\exp(-\Phi(x))$ with the vector of SEM weights $\boldsymbol{w}$. The eigenvectors are orthogonal with respect to the mass matrix: ${\boldsymbol{\phi_i}}^T\boldsymbol{W}_0\boldsymbol{\phi_j} = \delta_{ij}$, where $\boldsymbol{W}_0$ is a diagonal matrix with diagonal entries equal to the elementwise product of $\exp(-\Phi(x))$ with the vector of SEM weights $\boldsymbol{w}$.

After finding the eigenvalues $\lambda_{0,i}$ and eigenvectors $\phi_{0,i}(x)$ of the problem Eq. (9), we obtain the solution of the eigenvalue problem Eq. (8) by scaling back the eigenvectors: $\Psi_{0,i}(x) = \phi_{0,i}(x)\exp(-\Phi(x)/2)$, and the eigenvalues stay unchanged. It is convenient to represent the eigenvalues as a single vector $\boldsymbol{\lambda}_0 = \{\lambda_{0,i}\}$, and the eigenvectors as a transformation matrix that contains each eigenvector as a column $\boldsymbol{Q}_0 = \{\boldsymbol{\Psi}_0\}$. The scaling $\Psi_{0,i}(x) = \phi_{0,i}(x)\exp(-\Phi(x)/2)$ makes the new eigenvectors $\Psi(x)$ to be orthogonal with respect to the diagonal matrix of the SEM weights $\boldsymbol{W} = \mathrm{diag}(\boldsymbol{w})$, so that $\boldsymbol{Q}_0^T\boldsymbol{W}\boldsymbol{Q}_0 = \boldsymbol{I}$.

Next, we find the eigenvalues and eigenvectors of the operator $\boldsymbol{\mathcal{H}}$:

$$\boldsymbol{\mathcal{H}}\Psi(x) = \lambda\Psi(x). \tag{10}$$

To this end, we rewrite Eq. (10) in the basis of operator $\boldsymbol{\mathcal{H}}_0$. By using the definition $\boldsymbol{\mathcal{H}} = \boldsymbol{\mathcal{H}}_0 + \boldsymbol{\mathcal{H}}_I$, Eq. (10) in the basis of operator $\boldsymbol{\mathcal{H}}_0$ reads:

$$(\boldsymbol{\lambda}_0 \boldsymbol{I} + \boldsymbol{Q}_0^T \boldsymbol{W} \boldsymbol{F} \boldsymbol{Q}_0)\hat{\boldsymbol{\Psi}} = \lambda \hat{\boldsymbol{\Psi}}, \tag{11}$$

where $\boldsymbol{F}$ is the sum of tuning functions of all neurons $\sum_{k=1}^{M} f_k(x)$ discretized on the SEM grid into a diagonal matrix. After obtaining the matrix of eigenvectors $\hat{\boldsymbol{Q}}$, which is a transformation matrix from the basis of operator $\boldsymbol{\mathcal{H}}_0$ to the basis of $\boldsymbol{\mathcal{H}}$, we find the solution of the original problem as $\boldsymbol{Q} = \boldsymbol{Q}_0 \hat{\boldsymbol{Q}}$. The matrix $\boldsymbol{Q}$ contains the eigenvectors $\boldsymbol{\Psi}$ that solve the problem Eq. (10). The eigenvalues of Eq. (10) are the same as for Eq. (11), since the eigenvalues are basis independent.

In summary, to find the time-dependent solution of Eq. (6), we solve the corresponding eigenvalue problem Eq. (10). This problem is solved by finding the first $N_v$ eigenvalues and eigenvectors of Eq. (9), then multiplying the eigenfunctions $\phi_0$ by $\exp(-\Phi(x)/2)$ to obtain the eigenfunctions $\Psi_0$ of the problem Eq. (8) that constitute the matrix $\boldsymbol{Q}_0$. Using these eigenvectors and the eigenvalues $\boldsymbol{\lambda}_0$, we then solve another eigenvalue problem Eq. (11), from which we obtain the eigenvalues $\boldsymbol{\lambda}$ and the eigenvectors $\hat{\boldsymbol{Q}}$. The eigenvalues $\boldsymbol{\lambda}$ and the eigenvectors $\boldsymbol{Q} = \boldsymbol{Q}_0 \hat{\boldsymbol{Q}}$ are then the solution of Eq. (10).

For numerical solution, we discretize all functions and the eigenvalue problems in the SEM grid [1]. Thus, all probability densities in Eq. (3) become vectors of size $N$ and all operators become matrices of size $N^2$, where $N$ is the number of grid points (we set $N = 449$ for all model fits). The scaled transition probability density $\rho(x_{t_i}|x_{t_{i-1}})$ is solution of Eq. (6) and can be expressed in the basis of operator $\boldsymbol{\mathcal{H}}$ as:

$$\boldsymbol{\rho}_{i,i-1} \equiv \boldsymbol{T}_i = \mathrm{diag}\left(\exp\left(-\boldsymbol{\lambda}(t_i - t_{i-1})\right)\right), \tag{12}$$

where $\boldsymbol{\rho}_{i,i-1}$ is the transition matrix over latent states between the times $t_{i-1}$ and $t_i$ of the adjacent spikes. When the scaled probability density of latent states $\boldsymbol{\rho}_j$ at time $t_j$ is multiplied by the transition matrix $\boldsymbol{\rho}_{j+1,j}$ between the times $t_j$ and $t_{j+1}$, the result is the scaled probability density $\boldsymbol{\rho}_{j+1}$ at time $t_{j+1}$, since the matrix-vector product marginalizes over the distribution of latent states at time $t_j$. Similarly, the emission matrix, which gives the probability of observing a spike from neuron $k$ conditioned on the latent state, is expressed in the basis of operator $\boldsymbol{\mathcal{H}}$ as:

$$\boldsymbol{E}_k = \boldsymbol{Q}^T \boldsymbol{W} \mathrm{diag}(\boldsymbol{f}_{\mathrm{k}}) \boldsymbol{Q}, \tag{13}$$

where $\boldsymbol{f}_k$ is a vector of the discretized tuning function $f_k(x)$. The matrix of the absorption operator $\boldsymbol{A} = \hat{\boldsymbol{\mathcal{H}}}_0$ in the $\boldsymbol{\mathcal{H}}$-basis is:

$$\boldsymbol{A} = \hat{\boldsymbol{Q}}^T \mathrm{diag}(\boldsymbol{\lambda}_0) \hat{\boldsymbol{Q}}, \tag{14}$$

where $\hat{\boldsymbol{Q}}$ are the eigenvectors of the problem Eq. (11).

We can rewrite Eqs. (2), (3) in the finite basis of operator $\boldsymbol{\mathcal{H}}$ to obtain the chain of vector-matrix multiplications:

$$\mathscr{L} = \boldsymbol{\rho}_0^T \boldsymbol{T}_1 \boldsymbol{E}_{k_1} \boldsymbol{T}_2 \boldsymbol{E}_{k_2} \cdots \boldsymbol{T}_{N+1} \boldsymbol{A} \boldsymbol{\beta}_{N+2}. \tag{15}$$

Here $\boldsymbol{\beta}_{N+2}$ is a column vector used to integrate the likelihood over the final state $x_{t_E}$. This vector is equal to $\boldsymbol{\beta}_{N+2} = \boldsymbol{\rho}_{\mathrm{eq}} \boldsymbol{w}$, where $\boldsymbol{\rho}_{\mathrm{eq}}$ is a discretized vector of $\rho_{\mathrm{eq}}(x) = \sqrt{p_{\mathrm{eq}}} = \exp(-\Phi(x)/2)$ that scales $\rho(x,t)$ back into $p(x,t)$ and sums it up with the vector of SEM weights to integrate over

$x_{t_E}$. In Eq. (15), the indices $k_1, k_2, \dots k_N$ refer to the index of a neuron emitting a spike at times $t_1, t_2, \dots t_N$, and the spike times are ordered across all neurons.

We evaluate Eq. (15) with a forward pass that calculates the chain from left to right:

$$\begin{aligned} &\boldsymbol{\alpha_0}^T = \boldsymbol{\rho_0}^T, \qquad \boldsymbol{\alpha_n}^T = \boldsymbol{\alpha_{n-1}}^T \boldsymbol{T}_n \boldsymbol{E}_{k_n},\, n = 1, 2 \dots N, \\ &\boldsymbol{\alpha_{N+1}}^T = \boldsymbol{\alpha_N}^T \boldsymbol{T}_{N+1}, \qquad \boldsymbol{\alpha_{N+2}}^T = \boldsymbol{\alpha_{N+1}}^T \boldsymbol{A}, \end{aligned} \tag{16}$$

with $\mathscr{L} = \boldsymbol{\alpha_{N+2}}^T \boldsymbol{\beta_{N+2}}$. We use a scaling algorithm similar to that for the Hidden Markov Models [3]. After calculating each $\boldsymbol{\alpha_n}$, we calculate $c_n = ||\boldsymbol{\alpha_n}||$ and divide each $\boldsymbol{\alpha_n}$ by its norm before calculating $\boldsymbol{\alpha_{n+1}}$. With the scaling algorithm, each $\boldsymbol{\alpha_n}$ has the norm equal to 1, and the likelihood is equal to the product of all coefficients $c$, that is, $\log \mathscr{L} = \sum c_n$.

#### 2.2 Analytical derivation of the likelihood derivatives

We have previously derived the expressions for the variational derivatives of the likelihood with respect to the force $F(x)$, the auxiliary function $F_0(x)$, and the noise magnitude D [1]:

$$\begin{aligned} \frac{\delta \mathscr{L}}{\delta F(x)} &= \sum_{ij} G_{ij} \frac{D}{2} e^{-\Phi(x)} \frac{d\left(\phi_i(x)\phi_j(x)\right)}{dx} + \frac{1}{2}\int_{-1}^{x} \left(\beta_0(s) p_0(s) e^{\Phi(s)/2} - \alpha_{N+2}(s) e^{-\Phi(s)/2}\right) ds, \\ \frac{\delta \mathscr{L}}{\delta F_0(x)} &= \int_{-1}^{x} p_0(s) \left(\mathscr{L} - e^{\Phi(s)/2}\beta_0(s)\right) ds, \\ \frac{\partial \mathscr{L}}{\partial D} &= -\int_{-1}^{1} dx e^{-\Phi(x)} \sum_{ij} G_{ij} \frac{d\phi_i(x)}{dx}\frac{d\phi_j(x)}{dx}. \end{aligned} \tag{17}$$

Here the matrix $\boldsymbol{G}$ is computed with a forward-backward algorithm [3]. First, we perform the forward pass using Eq. (16). Then, we perform the backward path via a series of matrix-vector multiplications:

$$\begin{aligned} &\boldsymbol{\beta}_{N+1} = \boldsymbol{A}\boldsymbol{\beta}_{N+2}, \\ &\boldsymbol{\beta}_n = \boldsymbol{E}_{k_{N+1}} \boldsymbol{T}_{N+1} \boldsymbol{\beta}_{n+1}, \qquad n = 1, 2 \dots N, \\ &\boldsymbol{\beta}_0 = \boldsymbol{T}_1 \boldsymbol{\beta}_1. \end{aligned} \tag{18}$$

We compute the matrix $\boldsymbol{G}$ during the backward pass:

$$G_{ij} = \sum_{\tau=0}^{N+1} \Gamma_{ij}^{\tau+1} \alpha_{\tau,i} \beta_{\tau+1,j}, \tag{19}$$

where the matrix $\Gamma_{ij}^{\tau}$ is equal to the negative identity matrix for $\tau = N+2$, $\boldsymbol{\Gamma}^{N+2} = -\boldsymbol{I}$, and otherwise:

$$\Gamma_{ij}^{\tau} = \int_0^{\Delta t_\tau} e^{-(\Delta t_\tau - u)\lambda_i} e^{-u\lambda_j} du = \begin{cases} \Delta t_\tau e^{-\lambda_i \Delta t_\tau}, & i = j, \\ \frac{e^{-\lambda_i \Delta t_\tau} - e^{-\lambda_j \Delta t_\tau}}{\lambda_j - \lambda_i}, & i \neq j, \end{cases} \tag{20}$$

where $\Delta t_\tau = t_\tau - t_{\tau-1}$.

In this work, we also perform the inference of tuning functions $f_k(x)$ via the auxiliary functions $F_k(x)$ and auxiliary variables $C_k$. To derive the analytical expressions for these derivatives, we follow similar steps as in Ref. [1]. We write the expression for the likelihood in terms of propagation and emission operators:

$$\mathscr{L}\left[Y(t)|\theta\right] = \left\langle \rho_0 \middle| e^{-\boldsymbol{\mathcal{H}}\Delta t_1} \boldsymbol{y}_{k_1} e^{-\boldsymbol{\mathcal{H}}\Delta t_2} \boldsymbol{y}_{k_2} ... e^{-\boldsymbol{\mathcal{H}}\Delta t_N} \boldsymbol{y}_{k_N} e^{-\hat{\boldsymbol{\mathcal{H}}}\Delta t_{N+1}} \boldsymbol{A} \middle| \beta_{N+2} \right\rangle, \tag{21}$$

where we use the bra-ket notation for the state vectors and operator matrices. This form is basis independent and allows for the analytical likelihood calculation. In this notation, row vectors, such as $\boldsymbol{\rho}_0^T$, correspond to bra states $\langle \rho_0|$, propagation matrices, such as $\boldsymbol{T}_i$,correspond to the exponential operators $\exp\left(-\boldsymbol{\mathcal{H}}(t_i - t_{i-1})\right)$, spike emission matrices correspond to the spike emission operators, and column vectors, such as $\boldsymbol{\beta}_{N+2}$, correspond to ket states $|\beta_{N+2}\rangle$.

First, we compute the derivative of the likelihood with respect to the tuning function $f_k(x)$ of neuron $k$. In Eq. (21), each of the propagation operators depends on $f_k(x)$. In addition, the emission operators for which $k_i = k$ also depend on $f_k(x)$. Using the formula for product of the derivatives, we obtain:

$$\frac{\delta \mathscr{L}}{\delta f_k(x)} = \sum_{i,j} \left[ \sum_{\tau=1}^{N+1} a_i(\tau-1) b_j(\tau) \frac{\delta \langle \Psi_i | e^{-\boldsymbol{\mathcal{H}}\Delta t_\tau} | \Psi_j \rangle}{\delta f_k(x)} + \sum_{\tau_2} \hat{a}_i(\tau_2 - 1) \hat{b}_j(\tau_2) \frac{\delta \langle \Psi_i | f_k(x) | \Psi_j \rangle}{\delta f_k(x)} \right], \tag{22}$$

where in the second sum, the index $\tau_2$ takes the values of indices corresponding to spikes of the neuron $k$. The quantities $a, b, \hat{a}, \hat{b}$ are obtained via two forward and backward passes:

$$\begin{aligned}
&\langle \alpha_0 | = \langle \rho_0 |, \qquad \langle \alpha_n | = \langle \alpha_{n-1} | e^{-\boldsymbol{\mathcal{H}}\Delta t_n} \boldsymbol{y}_{k_n},\ n = 1, 2 \ldots N, \\
&\langle \alpha_{N+1} | = \langle \alpha_N | e^{-\boldsymbol{\mathcal{H}}\Delta t_{N+1}}, \qquad \langle \alpha_{N+2} | = \langle \alpha_{N+1} | \boldsymbol{A}, \\
&|\beta_{N+1}\rangle = \boldsymbol{A} |\beta_{N+2}\rangle, \\
&|\beta_n\rangle = \boldsymbol{y}_{k_n} e^{-\boldsymbol{\mathcal{H}}\Delta t_{n+1}} |\beta_{n+1}\rangle,\ n = 1, 2, \ldots N, \qquad |\beta_0\rangle = e^{-\boldsymbol{\mathcal{H}}\Delta t_1} |\beta_1\rangle, \\
&a_i(\tau) = \langle \alpha_\tau | \Psi_i \rangle, \qquad b_j(\tau) = \langle \Psi_j | \beta_\tau \rangle.
\end{aligned} \tag{23}$$

$$\begin{aligned}
&\langle \hat{\alpha_0} | = \langle \rho_0 | e^{-\boldsymbol{\mathcal{H}}\Delta t_1}, \qquad \langle \hat{\alpha}_n | = \langle \hat{\alpha}_{n-1} | \boldsymbol{y}_{k_n} e^{-\boldsymbol{\mathcal{H}}\Delta t_{n+1}},\ n = 1, 2 \ldots N, \\
&|\hat{\beta}_N\rangle = e^{-\boldsymbol{\mathcal{H}}\Delta t_{N+1}} \boldsymbol{A} |\hat{\beta}_{N+2}\rangle, \qquad |\hat{\beta}_n\rangle = e^{-\boldsymbol{\mathcal{H}}\Delta t_{n+1}} \boldsymbol{y}_{k_n+1} |\hat{\beta}_{n+1}\rangle, \\
&\hat{a}_i(\tau) = \langle \hat{\alpha}_\tau | \Psi_i \rangle, \qquad \hat{b}_j(\tau) = \langle \Psi_j | \hat{\beta}_\tau \rangle.
\end{aligned} \tag{24}$$

In the discretized space, $\langle \alpha_\tau|$ correspond to the forward vectors $\boldsymbol{\alpha}_\tau$ in Eq. (16), and $|\beta_\tau\rangle$ correspond to the backward vectors $\boldsymbol{\beta}_\tau$ in Eq. (18) with components $\alpha_i(\tau)$ and $\beta_i(\tau)$, respectively.

Using the formula for the derivative of the exponential operator, we obtain[4]:

$$\frac{\delta \langle \Psi_i | e^{-\boldsymbol{\mathcal{H}}\Delta t_\tau} | \Psi_j \rangle}{\delta f_k(x)} = -\left\langle \Psi_i \middle| \frac{\delta \boldsymbol{\mathcal{H}}}{\delta f_k(x)} \middle| \Psi_j \right\rangle \Gamma^\tau_{i,j}, \tag{25}$$

where the matrix $\Gamma$ is defined in Eq. (20). Operator $\boldsymbol{\mathcal{H}}$ depends on $f_k(x)$ through the term $\boldsymbol{\mathcal{H}}_I = \sum_{k=1}^M f_k(x)$, and we compute its variational derivative using the Euler-Lagrange equation:

$$\left\langle \Psi_i \middle| \frac{\partial \boldsymbol{\mathcal{H}}}{\partial f_k(x)} \middle| \Psi_j \right\rangle = \Psi_i(x) \Psi_j(x). \tag{26}$$

As a result, we obtain for the first term on the right hand side of Eq. (22):

$$\sum_{\tau=1}^{N+1} a_i(\tau-1)b_j(\tau)\frac{\delta\langle\Psi_i|e^{-\boldsymbol{\mathcal{H}}\Delta t_\tau}|\Psi_j\rangle}{\delta fr_k} = -\Psi_i(x)\Psi_j(x)\hat{G}_{ij}, \tag{27}$$

where

$$\hat{G}_{ij} = \sum_{\tau=0}^{N}\Gamma_{ij}^{\tau+1}\alpha_{\tau,i}\beta_{\tau+1,j}. \tag{28}$$

Note that $\hat{G}_{ij}$ is the same as $G_{ij}$ defined in Eq. (19) except for the last term, since the absorption operator does not depend on the tuning function and hence does not contribute to the derivative. We similarly evaluate the second term on the right hand side of Eq. (22) and obtain the likelihood derivative with respect to the tuning function:

$$\frac{\delta\mathscr{L}}{\delta f_k(x)} = \sum_{ij}\Psi_i(x)\Psi_j(x)(G_{ij}^{(k)} - \hat{G}_{ij}), \tag{29}$$

with the matrix $G_{ij}^{(k)}$ is defined as:

$$G_{ij}^{(k)} = \sum_{\tau_2}\hat{\alpha}_{\tau_2,i}\hat{\beta}_{\tau_2+1,j}, \tag{30}$$

where $\tau_2$ runs through all time points where neuron $k$ spiked.

Next, we compute the likelihood derivatives with respect to the auxiliary function $F_k(x)$ and the auxiliary variable $C_k$ used in the optimization. We note that

$$\frac{\partial f_k(x)}{\partial C_k} = \frac{f_k(x)}{C_k}. \tag{31}$$

To evaluate the derivative of $f_k(x)$ with respect to $F_k(x)$, we introduce another auxiliary function $R_k(x) = \int_{-1}^{1} F_k(x')H(x-x')dx' = \int_{-1}^{x} F_k(x')dx'$, where $H(s)$ is the Heaviside step function. With this substitution, $f_k(x) = C_k\exp(R_k(x))$, hence:

$$\begin{aligned}\frac{\delta R_k(s')}{\delta F_k(s)} &= H(s'-s),\\ \frac{\delta f_k(x)}{\delta R_k(s')} &= C_k e^{R_k(x)}\delta(s'-x).\end{aligned} \tag{32}$$

Using this results, we obtain:

$$\frac{\delta f_k(x)}{\delta F_k(s)} = \int_{-1}^{1}\frac{\delta f_k(x)}{\delta R_k(s')}\frac{\delta R_k(s')}{\delta F_k(s)}ds' = C_k e^{R_k(x)}\int_{-1}^{1}\delta(s'-x)H(s'-s)ds' = f_k(x)H(x-s). \tag{33}$$

Using the results in Eqs. (29), (33), we obtain the final expressions:

$$\begin{aligned}\frac{\delta\mathscr{L}}{\delta F_k(x)} = \int_{-1}^{1}\frac{\delta\mathscr{L}}{\delta f_k(s)}\frac{\delta f_k(s)}{\delta F_k(x)}ds &= \int_{-1}^{1}\frac{\delta\mathscr{L}}{\delta f_k(s)}f_k(s)H(s-x)ds =\\ &= \int_{-1}^{1}\frac{\delta\mathscr{L}}{\delta f_k(s)}f_k(s)ds - \int_{-1}^{x}\frac{\delta\mathscr{L}}{\delta f_k(s)}f_k(s)ds,\end{aligned} \tag{34}$$

and for the derivative with respect to the constants $C_k$ we obtain:

$$\frac{\partial \mathscr{L}}{\partial C_k} = \int_{-1}^{1} \frac{\delta \mathscr{L}}{\delta f_k(s)} \frac{\partial f_k(s)}{\partial C_k} ds = \frac{1}{C_k} \int_{-1}^{1} \frac{\delta \mathscr{L}}{\delta f_k(s)} f_k(s) ds. \tag{35}$$

In summary, we compute the derivatives of the likelihood with respect to $F$, $F_0$ and $D$ using Eq. (17) derived in Ref. [1]. The derivatives with respect to $F_k(x)$ and $C_k$ for each neuron $k$ are computed using Eqs. (34), (35) and (29).

#### 2.3 Maximum likelihood optimization with ADAM algorithm

We optimized the model likelihood using a modified ADAM gradient-descent algorithm combined with line searches. The ADAM update on iteration $t$ is:

$$\begin{aligned} m_t &= \beta_1 m_{t-1} + (1-\beta_1) g_t, \\ \hat{m}_t &= m_t/(1-\beta_1^t), \\ v_t &= \beta_2 v_{t-1} + (1-\beta_2)||g_t||^2, \\ \hat{v}_t &= v_t/(1-\beta_2^t), \\ \theta_t &= \theta_{t-1} - \alpha \frac{\hat{m}_t}{\sqrt{\hat{v}_t} + \epsilon}, \end{aligned} \tag{36}$$

where $\alpha$, $\beta_1$, $\beta_2$, $\epsilon$ are hyperparameters, $m_t$ is a running average of the gradient $g_t$ on iteration $t$, $v_t$ is a running average of the gradient's $L^2$ squared norm, $\hat{m}_t$ and $\hat{v}_t$ are computed from $m_t$ and $v_t$ to correct the initialization bias (at zero iteration $m_0 = v_0 = 0$). We apply Eqs. (36) independently to each of the model's components $F(x), F_0(x), F_i(x), D, C_i$. The main difference between the original ADAM algorithm and our version is that we optimize over the space of continuous functions $F(x)$, $F_0(x)$, and $\{F_i(x)\}$ and not independent discrete parameters, therefore we scale their gradients by the average function's $L^2$-norm defined as $\|\phi(x)\|_2 = \int_x |\phi(x)|^2 dx$.

On each epoch of ADAM, we randomly group the trials into 20 identical trial batches. Each epoch thus consisted of 20 iterations. We then compute ADAM quantities and update each of the model components using Eqs. (36) on each batch of trials. In addition, on a subset of epochs, we performed line searches for the scalar parameters $D$ and each $C_i$ using L-BFGS-B method from scipy.optimize.minimize toolbox. Since a line search is computationally expensive, we perform only 30 line searches spaced logarithmically over the 5,000 epochs range, such that most line searches are concentrated at early epochs.

To perform shared optimization across four stimulus conditions, we randomly split trials for each of the four conditions into 20 batches of equal size, resulting in 80 batches total. We randomly permuted the order of these 80 batches, but all trials in one batch come from the same condition. For each trial batch, we calculated the gradients and updated the model components Eqs. 36. In this case, we optimized four potential functions $\Phi_1(x), \Phi_2(x), \Phi_3(x), \Phi_4(x)$, each of which was updated only on trial batches from the corresponding condition. Other parameters, including $D$, $F_i(x)$, $C_i$ and $F_0(x)$ were the same for all conditions, and thus were updated 80 times per epoch on all trial batches.

For bootstrapping, we first randomly divided our dataset into two equally sized data samples, and then sampled the trials randomly with replacement from each of the data samples. Thus, our bootstrap samples from two data halves did not intersect with each other but both of them contained repeated trials.

#### 2.4 Model selection

We used the model selection procedure based on feature consistency that we developed in Ref. [1]. First, we introduce feature complexity of a model $\mathcal{M} = -S[\Phi(x), D, p_0(x); \Phi^R(x), D^R, p_0^R(x)]$ defined as a negative trajectory entropy. The trajectory entropy is defined as a negative Kullback-Leibler (KL) divergence between the distributions $P[\mathcal{X}(t)]$ and $Q[\mathcal{X}(t)]$ [5]:

$$S[\Phi(x), D, p_0(x); \Phi^R(x), D^R, p_0^R(x)] = -\int_0^{t_{\text{obs}}} \mathscr{D}\mathcal{X}(t) P\left[\mathcal{X}(t)\right] \ln \frac{P\left[\mathcal{X}(t)\right]}{Q\left[\mathcal{X}(t)\right]}. \tag{37}$$

$P[\mathcal{X}(t)]$ is the distribution of trajectories in the model of interest with Langevin parameters $\{\Phi(x), D, p_0(x)\}$, and $Q[\mathcal{X}(t)]$ is the distribution of trajectories in the reference model with Langevin parameters $\{\Phi^R(x), D^R, p_0^R(x)\}$. The path integral is performed over all possible trajectories $\mathcal{X}(t)$. The reference model is a free diffusion with zero driving force (i.e. constant potential $\Phi^R(x) = \text{const}$), uniform $p_0^R(x)$, and the same diffusion coefficient $D$ as in the model of interest. Intuitively, feature complexity quantifies the number and prominence of features in the model: the greater $\mathcal{M}$ the more complex trajectories the model generates.

We previously derived an analytical expression for the trajectory entropy through the Langevin parameters [1]:

$$S[\Phi(x), D, p_0(x); \Phi^R(x), D, p_0^R(x)] = -\int_x dx p_0(x) \ln \frac{p_0(x)}{p_0^R(x)} - \frac{D}{4}\int_0^\infty dt \int_x dx F^2(x) p(x,t). \tag{38}$$

To evaluate Eq. (38) numerically, we integrate the first term using the Gaussian quadrature on a Gauss-Legendre-Lobatto (GLL) grid:

$$\int dx p_0(x) \ln \frac{p_0(x)}{p_0^R(x)} = \sum_k p_0(x_k) \ln \frac{p_0(x_k)}{p_0^R(x_k)} w_k, \tag{39}$$

where $w_k$ are the GLL integration weights. The second term can be simplified by switching into the basis of operator $\boldsymbol{\mathcal{H}}$ and evaluating the integral over time analytically [1]:

$$\int_0^\infty dt \int dx F^2(x) p(x,t) = \sum_k \frac{\rho_{0,k}(F^2 \exp[-\Phi/2])_k}{\lambda_k}. \tag{40}$$

Here $k$ indexes the elements of the vectors in the basis of operator $\boldsymbol{\mathcal{H}}$, $\rho_0(x) = p_0(x)/\sqrt{p_{\text{eq}}(x)}$, and $\lambda_k$ are the eigenvalues of operator $\boldsymbol{\mathcal{H}}$. Thus, Eqs. (38–40) allow us to compute feature complexity from the Langevin parameters $\Phi(x)$, $p_0(x)$, and $D$.

We split our full dataset into two halves $\mathcal{D}_1$ and $\mathcal{D}_2$ and optimize the model on each data split independently. As a result, we obtain two sequences of models $\theta_{1,n}$ and $\theta_{2,n}$, where $n = 1, 2, \ldots 5,000$ is the epoch number. To reduce the amount of computations needed for model selection, we then subsample these models on a logarithmic scale by choosing 400 models indexed by $m$ where $m \propto \log n$. For each of the sub sampled models we then calculate feature complexity $\mathcal{M}(\Phi(x), p_0(x), D)$ and obtain two sequences $\mathcal{M}_{1,m}$, and $\mathcal{M}_{2,m}$ for $m = 1, 2, ..., 400$. For each level of feature complexity, we compare two models optimized on $\mathcal{D}_1$ and $\mathcal{D}_2$. We quantify the consistency of features between models using Jansen-Shannon divergence between their time-dependent probability distributions [1]:

$$D_{\text{JS}} = \int_0^\infty \text{JSD}\left(\hat{p}^1(x,t) || \hat{p}^2(x,t)\right) dt. \tag{41}$$

Here $\hat{p}^1(x,t)$ and $\hat{p}^2(x,t)$ are the time-dependent probability densities of latent states generated by the two models. The Jason-Shanon divergence between two distributions is computed as:

$$\begin{aligned}\mathrm{JSD}(\hat{p}^1(x)||\hat{p}^2(x)) = \frac{1}{2}\left(\int \hat{p}^1(x)\log\frac{2\hat{p}^1(x)}{\hat{p}^1(x)+\hat{p}^2(x)}dx + \int \hat{p}^2(x)\log\frac{2\hat{p}^2(x)}{\hat{p}^1(x)+\hat{p}^2(x)}dx\right)+ \\ +\frac{1}{2}\left(I_1\log\frac{2I_1}{I_1+I_2} + I_2\log\frac{2I_2}{I_1+I_2}\right),\end{aligned} \quad (42)$$

where $I_{1,2} = 1 - \int \hat{p}^{1,2}(x)dx$. We compute $D_{\mathrm{JS}}$ by a forward Euler time discretization of Eq. (41), where for each time step the integral Eq. (42) is computed with the GLL integration weights $w_k$.

At early optimization epochs the models of similar feature complexities optimized on two different datasamples agree with each other. At some feature complexity level $\mathcal{M}^*$ the two models start to disagree due to overfitting to noise which produces features that are different for two independent data samples. We numerically identify $\mathcal{M}^*$ at the point where $D_{\mathrm{JS}}$ exceeds a threshold $D_{\mathrm{thres}} = 0.0015$.

Since feature complexities do not match exactly between the two model sets due to nuances in the data, we need to allow for some slack in feature complexity when comparing models[1]. Accordingly, for an iteration $m_1$ on the data split $\mathcal{D}_1$, we first find iteration $\hat{m}_2$ on $\mathcal{D}_2$ which minimizes $|\mathcal{M}_{1,m_1} - \mathcal{M}_{2,\hat{m}_2}|$. Next, we swipe $m_2 = \hat{m}_2 - 5, \hat{m}_2 - 4, \ldots \hat{m}_2 + 5$, and select index $m_2$ that minimizes $D_{\mathrm{JS}}$ between models $m_1$ and $m_2$, and set $D_{\mathrm{JS}}(\mathcal{M}_{1,m_1})$ to this minimum value. We repeat this procedure for different epochs to obtain the dependence $D_{\mathrm{JS}}(\mathcal{M})$ to which we apply the threshold $D_{\mathrm{JS,thres}}$. To reduce the computational cost, we evaluate $D_{\mathrm{JS}}(\mathcal{M})$ on a subset of 400 epochs spaced logarithmically over the 5,000 optimization epochs range, so that earlier epochs are sampled more densely.

#### 2.5 Viterbi algorithm

We generalized the max-sum Viterbi algorithm with backtracking [3] to our case of continuous-space continuous-time latent dynamical system. The goal of the Viterbi algorithm is to find a path $X(t)$ that maximizes the joint probability density $P(X(t), Y(t))$ in a particular trial. The discretized path $X(t) = \{x_{t_0}, x_{t_1}, \ldots, x_{t_N}, x_{t_E}\}$ is the sequence of states at the trial start time, each of the spike times, and the trial end time. We describe the Viterbi algorithm for a single-neuron model, and the extension to a population model is straightforward.

We maximize the logarithm of the joint probability density:

$$\log P(X(t), Y(t)) = \log p(x_{t_0}) + \log p(x_{t_1}|x_{t_0}) + \log p(y|x_{t_1}) + \log p(x_{t_2}|x_{t_1}) \cdots + \log p(A|x_{t_E}). \quad (43)$$

In Eq (43), only the first two terms depend on $x_{t_0}$, and their sum is the joint probability density $\log p(x_{t_1}, x_{t_0})$. To compute the joint probability density in the discretized latent domain with $N$ grid points, we transform the vector $\log p(x_{t_0})$ into an $N \times N$ matrix with the same rows and sum it with the $N \times N$ matrix representing $\log p(x_{t_1}|x_{t_0})$. This operation is similar to the likelihood computation but without integration over $x_{t_0}$. We then take the maximum over $x_{t_0}$ to obtain a vector $A_1(x_{t_1})$:

$$A_1(x_{t_1}) = \max_{x_{t_0}}\{\log p(x_{t_0}) + \log p(x_{t_1}|x_{t_0})\}. \quad (44)$$

In addition, we compute another vector $B_1(x_{t_1})$ that for each $x_{t_1}$ returns the index of $x_{t_0}$ (1 to $N$) that maximizes this expression:

$$B_1(x_{t_1}) = \underset{x_{t_0}}{\operatorname{argmax}} \{\log p(x_{t_0}) + \log p(x_{t_1}|x_{t_0})\}. \tag{45}$$

Similarly, we continue evaluating the terms in Eq. (43): we take the vector $A_1(x_{t_1})$, transform it into $N \times N$ matrix, add it to the terms that depend on $x_{t_1}$: $\log p(y|x_{t_1}) + \log p(x_{t_2}|x_{t_1})$, and compute the vectors $A_2(x_{t_2})$ and $B_2(x_{t_2})$ by taking the maximum and argmaximum over $x_{t_1}$, and so on. In the end of this procedure, we have computed the whole chain and we find the maximized joint probability density:

$$\max_{X(t)} [\log P(X(t), Y(t))] = \max_{x_{t_E}} [A_{N+1}(x_{t_E}) + \log p(A|x_{t_E})] . \tag{46}$$

To evaluate the most probable sequence of latent states, we then perform backtracking. Using the value of $x^*_{t_E}$ that maximizes Eq. (46), we find the latent state at the last spike time by retrieving its index: $x^*_{t_N} = x[B_{N+1}(x^*_{t_E})]$, and then continue backtracking until we recover the whole sequence of states that maximizes $P(X(t), Y(t))$.

For our case of absorbing boundary conditions, we introduce an additional modification to the Viterbi algorithm forcing the latent trajectory to reach one of the boundaries for the first time exactly at the trial end time. First, after adding each of the latent propagation terms, we zero out all boundary elements in the $N \times N$ matrix before the max operation Eq. (44), which prevents the latent trajectory reaching a boundary before the trial end. Second, on the last step Eq. (46), we restrict $x^*_{t_E}$ to be either the left or right boundary, $x^*_{t_E} = -1$ or $x^*_{t_E} = 1$, depending on which of the two values maximizes $\log P(X(t), Y(t))$ in Eq. (46).

### 3 Supplementary References

### References

[1] Genkin, M., Hughes, O. & Engel, T. A. Learning non-stationary Langevin dynamics from stochastic observations of latent trajectories. *Nat. Commun.* **12**, 5986 (2021).

[2] Genkin, M. & Engel, T. A. Moving beyond generalization to accurate interpretation of flexible models. *Nat. Mach. Intell.* **2**, 674–683 (2020).

[3] Bishop. *Pattern Recognition and Machine Learning* (Springer, New York, 2007).

[4] Wilcox, R. M. Exponential operators and parameter differentiation in quantum physics. *J. Math. Phys.* **8**, 962–& (1967).

[5] Haas, K. R., Yang, H. & Chu, J.-W. Trajectory entropy of continuous stochastic processes at equilibrium. *J. Phys. Chem. Lett.* **5**, 999 – 1003 (2014).
